# An image-computable spatio-chromatic receptive field model of the midget retinal ganglion mosaic across the retina

**DOI:** 10.1101/2025.10.20.683437

**Authors:** Nicolas P. Cottaris, Brian A. Wandell, David H. Brainard

## Abstract

Accurate image-computable models of retinal ganglion cell (RGC) mosaics across the retina do not currently exist. Here, we deploy a novel computational framework which synthesizes mosaics of linear spatio-chromatic receptive fields (RFs) of ON midget RGCs (mRGCs) by integrating published anatomical, physiological, and optical quality measurements. We use the synthesized mRGC mosaics to simulate both *in vivo* and *in vitro* physiological experiments and demonstrate the model’s consistency with published data. The model enables computation of how visual performance is shaped by the representation of visual information provided by the linear spatiochromatic processing stage of midget RGCs. The developed computational framework carefully accounts for the effect of physiological optics on mRGC responses, enables comparison of *in vivo* and *in vitro* data, and allows exploration of how different assumptions about RF organization, such as selectivity for the type of cones pooled by the RF center mechanism, affect physiological responses and psychophysical performance. The open-source and freely available implementation provides a platform for understanding how the linear spatiochromatic receptive field representation of the mRGCs shapes visual performance, as well as a foundation for future work that incorporates response nonlinearities, temporal filtering, and extends to additional RGC mosaics.

## 1 Introduction

An important aim in computational visual neuroscience is to create accurate computer simulations of how neurons in the visual pathways encode and respond to visual scenes. These simulations, often called digital twins, are a quantitative description of the visual system. They enable links between the neural representation and perception and provide a tool for evaluating the effects of blinding disease and its treatment.

Over the last ten years we have built an open-source software platform, ISETBio (Image Systems Engineering Tools for Biology) [1], which serves as a digital twin for the initial stages of the human visual system. Previously, we described how ISETBio models (a) the formation of the retinal image, (b) the excitation of the cone photoreceptors, (c) phototransduction, and (d) fixational eye movements [2–4]. We and others have employed ISETBio to model human vision, including sensitivity to spatial contrast [2, 3], the impact of chromatic aberration on acuity [5], the encoding of information from natural images captured by cones [6], the effects of optics and cone density across the visual field on performance [7], and the influence of initial visual signals on tasks like judging surface properties and lighting [8, 9]. We also used ISETBio to help interpret experimental measurements of retinal ganglion cells [10].

Here, we describe an extension of ISETBio to model the mosaic of a class of retinal ganglion cells (RGCs), the midget RGC mosaic. RGCs are the only pathway for information transmission from the retina to the brain, and their properties surely impact visual performance on many tasks. The signals sent from the one million RGC axons that form the human optic nerve represent the signals from roughly 6.5 million cones and 110 million rods [11, 12]. Of these RGCs, mRGCs are a particularly important subtype, comprising 80% of the perifoveal RGCs and 45% of the peripherial RGCs. In the very central fovea, it has been estimated that the mRGCs are 95% of the RGC population [13].

The role of the mRGCs in limiting spatial and color vision is still debated [14]. Simulation of performance using image computable models of the mRGC mosaic offers a powerful tool for understanding the visual information encoded by these cells, especially because they are very hard to measure and isolate experimentally. We have four primary goals for this human retina model.

First, the model must distinguish the contributions of the eye’s optics and photoreceptors from the subsequent post-receptoral retinal circuitry. This separation is crucial for incorporating key physiological measurements, some of which are made *in vitro* without the eye’s optics. Failing to isolate the optical effects would prevent us from using this vital collection of data.

Second, the model must capture responses across a large portion of central retina. This is important because we and others are interested in how the retinal representation shapes performance not just in the fovea but also for peripheral viewing. Third, the model must integrate diverse data types, including optical, anatomical, and physiological measurements. A comprehensive formulation is necessary because retinal ganglion cell (RGC) responses are shaped by all three of these factors.

Fourth, we aim for an extensible framework. The current implementation uses a linear spatiochromatic receptive field, which serves as a good initial approximation. The framework is designed to incorporate future extensions—such as response nonlinearities and additional RGC classes—to improve the model’s accuracy over time.

The following points describe how our implementation achieves these goals.

1. *Separating representations*. Our mRGC model operates on the cone mosaic signals. This design isolates the post-receptoral circuitry (cone-to-mRGC), which is the pathway measured in *in vitro* experiments where the eye’s optics are removed [15, 16]. This separation is also valuable for interpreting experiments that use adaptive optics to eliminate optical blur [10]. While the components are separable, our implementation integrates the optics, cone sampling, and mRGC circuitry into a complete, image-computable pipeline. This full pathway allows us to simulate the transformation of a visual stimulus into an mRGC response, matching the conditions of *in vivo* measurements [17–19] and enabling predictions of human performance under natural viewing conditions.
2. *Representation across the visual field*. Visual performance varies across the visual field, and a key contribution of our model is that it allows computation of the mRGC representation continuously across the retina from the fovea to 30°, along any meridian. Achieving this goal required implementation of novel algorithms for synthesizing mRGC RF mosaics.
3. *Multiple data types*. By explicitly representing different biological stages, our model enables algorithms that combine anatomical, physiological, and optical data. Incorporation of multiple types of measurements from the literature is critical because at present no one type of data sufficiently constrains mRGC properties across the visual field.
4. *Extensible*. The current implementation is a linear spatial pooling model, a useful approximation for stimuli with modest contrast. The software’s modular design provides a foundation for future extensions. We can incorporate known nonlinear properties that shape mRGC responses, including phototransduction effects [20]; spatial and static nonlinearities, which often differ between ON and OFF pathways [21–24]; temporal dynamics [25]; and response noise [26]. Furthermore, the mRGC model is a suitable base for developing models of other RGC types, such as parasol and bistratified cells [27].

### 1.1 Model overview

Fig. 1 provides a model overview. Computation begins with the image spectral radiance, such as produced by a calibrated monitor. A model of the human optics (including chromatic aberrations) and spectral filtering by the lens is used to compute the retinal irradiance. Retinal irradiance is spectrally filtered by the macular pigment and then spatially and spectrally sampled by the cone photoreceptor mosaic. The parameters of the optics, macular pigment and cone mosaic all vary across the visual field, according to measurements in the literature [2].

**Fig. 1.**
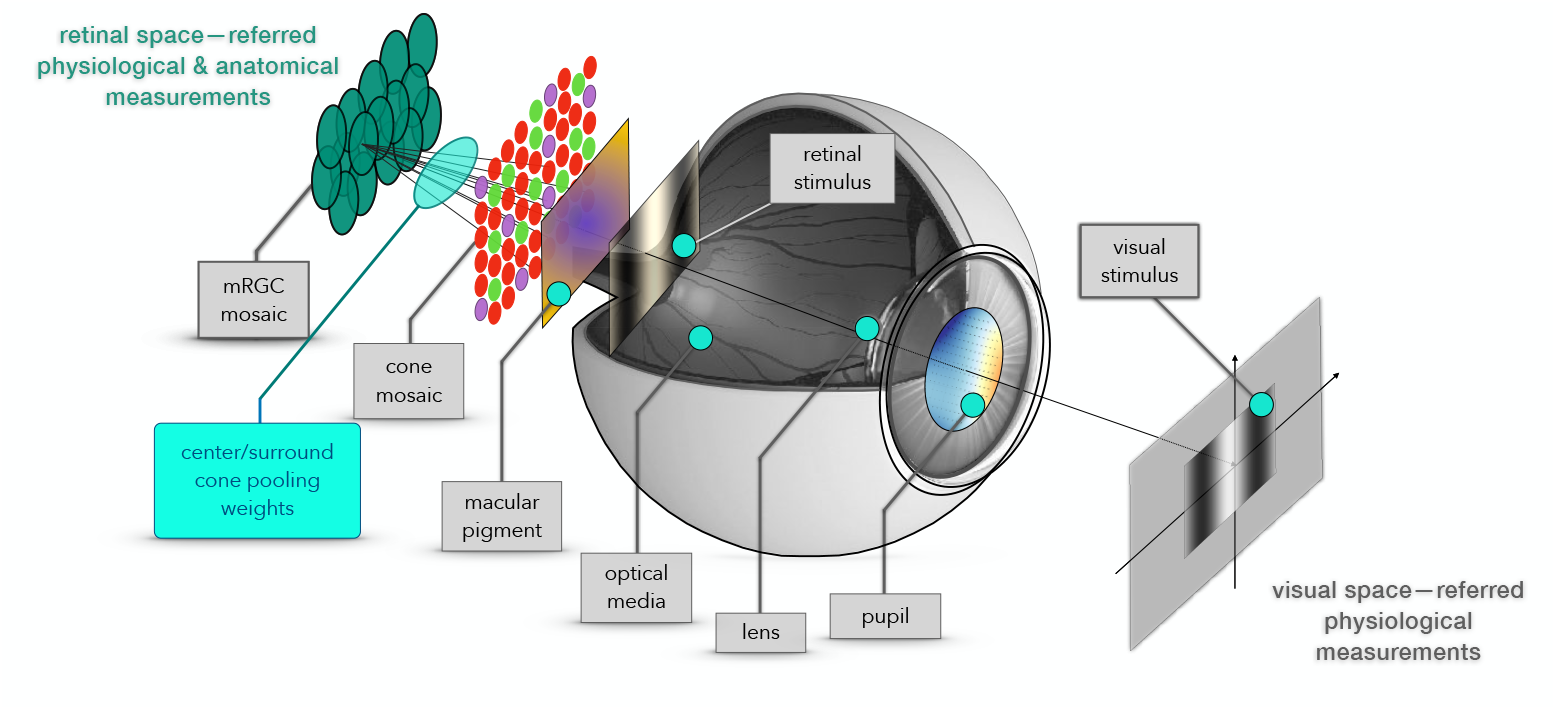
Model overview. The extant ISETBio model computes the mosaic of cone excitations. The model mRGCs are obtained by connecting their RFs to the cone mosaic. The connectivity matrix is constrained by anatomy and optimized through forward simulation of physiological measurements, so that the synthetic mRGCs are consistent with data across the visual field.

The mRGC mosaic extension is composed of spatial receptive fields (RFs) whose center and surround responses are weighted sums of signals from the cone mosaic.

The wiring between the input cone mosaic and the mRGC mosaic is initially determined based on anatomical constraints, such as cone and mRGC densities, and is subsequently refined using optimization algorithms that align the model’s spatial RF properties with physiological measurements.

A key challenge is the scarcity of *in vitro* physiological data across the visual field which could be used to directly determine the wiring between the two mosaics. To address this, our algorithm primarily leverages more widely available *in vivo* data for the optimization, while validating the model against *in vitro* data where it exists. The resulting model is simultaneously consistent with cone light encoding, anatomical properties (including those of mRGCs and H1 horizontal cells), and both *in vitro* and *in vivo* physiological data. This makes the model versatile for simulating visual stimulation under *in vivo, in vitro*, and adaptive optics paradigms.

The remainder of this paper is organized as follows.

- In section 2 we describe the model’s construction stages, including, how the mRGC receptive field lattice is generated from anatomical data (section 2.1), how cones get connected to the mRGC RF centers using anatomical and physiological constraints (section 2.2), and how cone connections to mRGC RF surrounds are derived by optimizing against *in vivo* data (section 2.3).
- In section 3 we present, validate, and discuss a first application of the model. Specifically, we illustrate examples of synthesized mRGC mosaics (section 3.1), confirm that the model mRGC spatial RFs are consistent with *in vivo* (section 3.2), and *in vitro* data (section 3.3), demonstrate the significant impact of physiological optics (section 3.4), how simpler Difference-of-Gaussians models can fail to capture the true surround pooling (section 3.5), and finally we present first applications of the developed model, which illustrate how the model can be used to estimate the contribution of the mRGC mosaic to spatiochromatic contrast sensitivity across the visual field (section 3.6).
- In section 4, we summarize our work, present some of the model’s applications, the model’s current limitations, and planned expansions.

## 2 Methods

The synthesis of mRGC RF mosaics occurs in three stages. In the first stage, we generate spatial lattices representing the RF centers of the mosaic of mRGCs and the position of cones of the cone mosaic that provides input to the mRGC mosaic. In the second stage, we connect the input cone mosaic to the RF centers of the mRGC mosaic. In the third stage, we connect the input cone mosaic to the RF surrounds of the mRGC mosaic.

### 2.1 Generating the spatial position lattice of mRGC RF centers (Stage 1)

We begin by generating a lattice that represents the (*x, y*) positions of mRGC RF centers. This process comprises three sub-stages illustrated in Fig. 2.

**Fig. 2.**
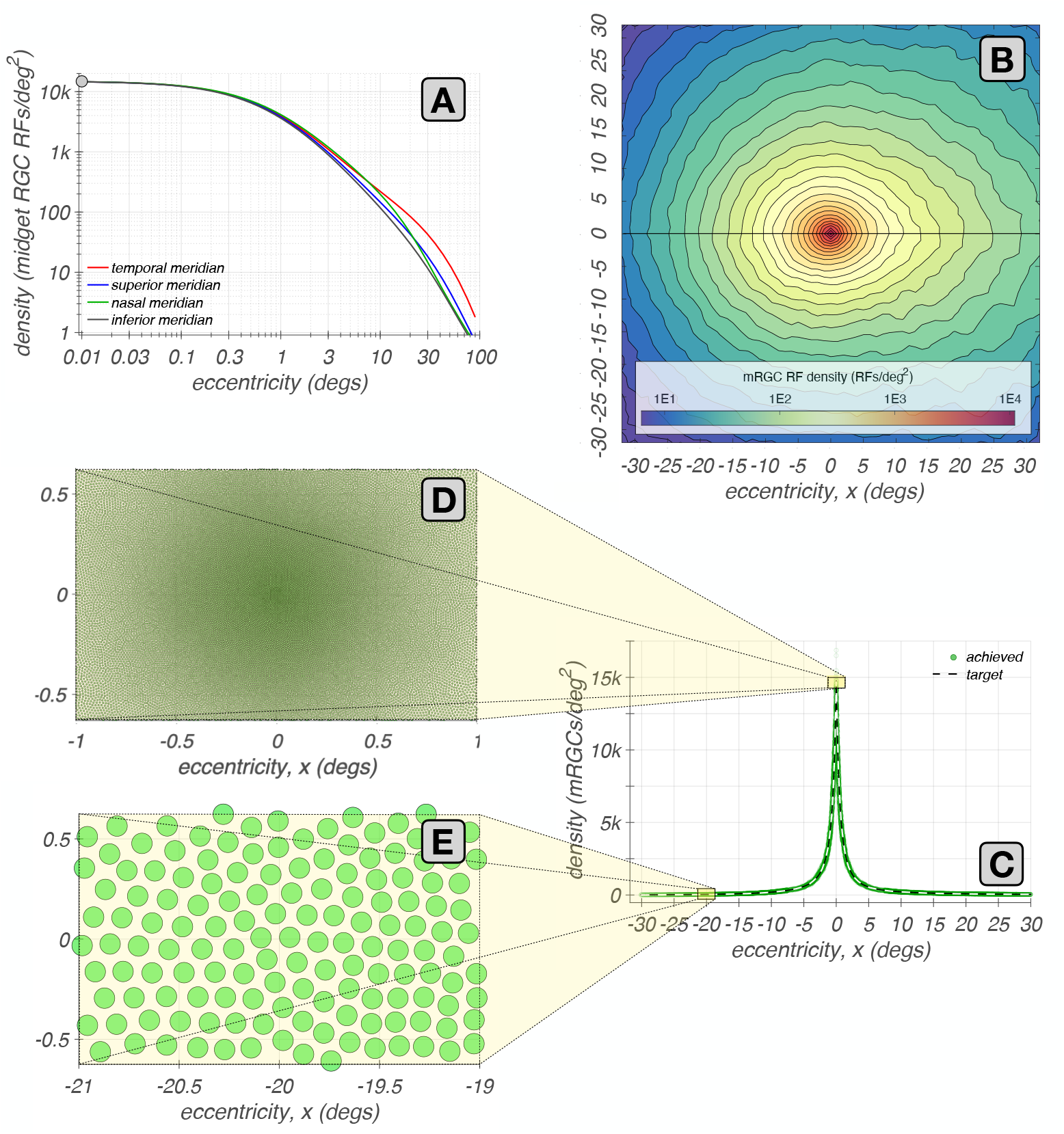
Eccentricity-varying mRGC RF position lattices. **A:** Meridian density functions of mRGC RFs [28]. **B:** Two-dimensional mRGC RF density map obtained by interpolating the four meridian density functions. **C:** Achieved and target densities of mRGC RF centers along the horizontal meridian (green disks and white dashed line). **D & E:** Examples of 2° *×* 1° mosaics of mRGC RF centers at eccentricities of 0° and 20°, respectively, along the temporal meridian.

- **Stage 1A**. We estimate the mRGC RF center densities along the four principal meridians (0°, 90°, 180°, and 270°). These estimates are based on human data [28, 29]. We take the ON-mRGC density to be half of the total mRGC density, ignoring the possible density differences between ON– and OFF–center mRGCs. The meridian functions are depicted in Fig. 2A.
- **Stage 1B**. We generate a continuous, two-dimensional map representing the mRGC RF density map, depicted in Fig. 2B. This map is created by linearly interpolating the meridian estimates, and it serves as a target for the algorithm in the next stage.
- **Stage 1C**. We synthesize a sampling lattice that represents the mRGC RF center positions. The lattice is created using the iterative algorithm that we introduced earlier [2] for generating cone mosaics, replacing the two-dimensional cone density map with the target mRGC RF density map. A typical lattice of mRGC RF positions is obtained after about 1,300 iterations and has a density that varies smoothly over space, matching the target density, as illustrated in Fig. 2C. Example patches of mRGC RF center mosaics synthesized at eccentricities of 0° and 20° along the temporal horizontal meridian, are depicted in Figs. 2D & E, respectively.

### 2.2 Connecting cones to mRGC RF centers (Stage 2)

The connections between cones and mRGC centers are constrained by (1) anatomical data across the retina, specifically, the ratio of densities of mRGC RF centers to cones [28], and (2) *in-vitro* physiological data from peripheral retina, that (a) indicate that, unlike OFF-center mRGCs, which draw indiscriminately from all three cone types [15, 30, 31], ON-center mRGCs draw only from L- and M-cones, and (b) quantify the degree of RF center overlap in neighboring mRGCs [32]. In the present version of the model, we only model ON-center mRGCs. The cone to mRGC RF center connectivity is established in 3 sub-stages, summarized here.

- **Stage 2A**. In the first substage, each L- and M-cone gets connected to a single mRGC RF center; single RF centers can receive input from more than one cone. S-cones are not connected because they do not contribute to ON-center mRGCs. This initial cone-to-RF center connectivity often results in inhomogeneities in the composition of neighboring mRGCs RF centers, which are dealt with in the nest stage.
- **Stage 2B**. This stage refines the center connections to establish a balance between the spectral purity and spatial compactness of the mRGC RF centers, which is quantified by a single parameter, *ϕ*. For the body of this work, all mRGC mosaics are generated by maximizing spatial compactness, but the option to maximize spectral purity allows testing of different scenarios where mRGC RF centers may be biased to some extent towards cone type selective pooling [15, 16].
- **Stage 2C** - Finally, the mutual exclusivity constraint enforced in stages 2A and 2B is lifted, and single cones are permitted to connect to multiple nearby mRGC RF centers. The extent of divergence varies with retinal eccentricity, being minimal in the fovea and increasing towards the periphery to match experimental observations [32]. This is done by varying the exponent of a supra-Gaussian distribution that describes the spatial weighting profile of cone connections to the RF centers, as described later.

The algorithms that implement each of these substages are described in detail in Section A. We illustrate Stage 2 by examining key properties of synthesized mRGC RF center mosaics at each of the three substages.

#### 2.2.1 Mosaics with convergent-only cone connections (stage 2A)

Example mosaics of RF centers synthesized at four eccentricities along the temporal horizontal meridian at the end of stage 2A are depicted in Fig. 3, where each green ellipse represents the spatial extent of the RF center of a single mRGC. For the foveal mosaic depicted in Fig. 3A, RF centers connect to just a single cone. Note how RF center sizes increase as we move towards parafoveal regions to the left and right sides of Fig. 3A. This is due to the continuously increasing, with eccentricity, cone aperture in the input cone mosaic. The empty regions in this foveal mRGC RF center mosaic correspond to the location of S-cones which are not pooled by the model.

**Fig. 3.**
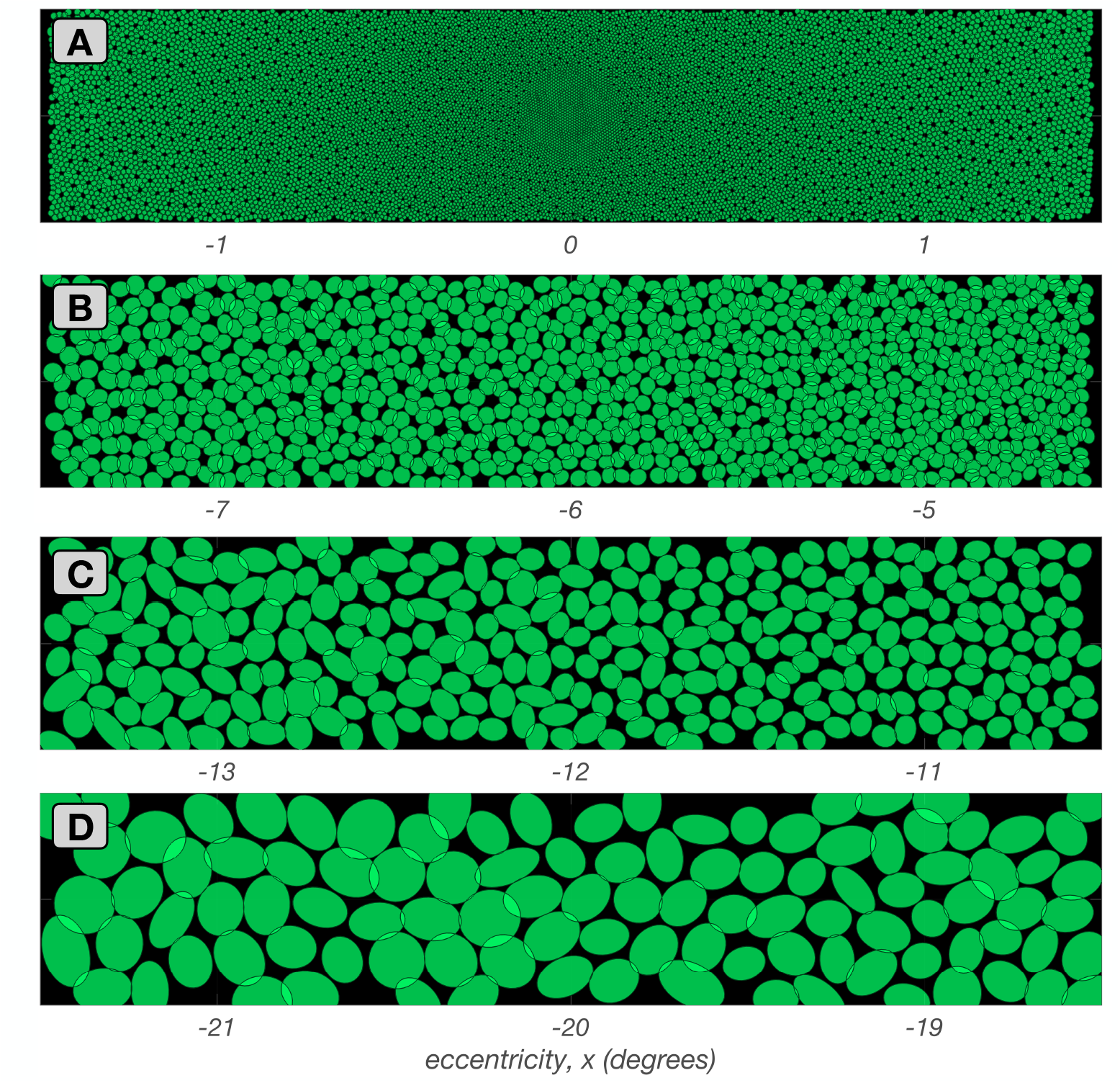
Stage 2A mRGC RF mosaics. Each panel shows a 3.0^*o*^ *×* 0.5^*o*^ mosaic of synthesized mRGC RF centers at a different visual field location from fovea to periphery. The green ellipses depict a spatial region that encompasses all cones pooled by single RF centers. **A:** Foveal mosaic, in which RF centers receive signals from a single L– or M–cone. **B:** Mosaic centered at 6.0^*o*^ along the temporal horizontal meridian, in which RF centers receive signals from 2–3 L/M–cones. **C:** Mosaic centered at 12.0^*o*^ along the temporal horizontal meridian, in which RF centers receive signals from 3–4 L/M–cones. **D:** Mosaic centered at 20.0^*o*^ along the temporal horizontal meridian, in which RF centers receive signals from 6–9 cones.

In the parafoveal mosaic depicted in Fig. 3B, RF centers mostly receive inputs from two cones, whereas in the more peripheral mosaics depicted in Figs 3C & 3D, RF centers connect to multiple cones. Note that the number of cones connecting to RF centers does not correspond precisely to RF center size, because cone aperture and inter-cone spacing both increase with eccentricity. At all eccentricities, however, mRGC RF center mosaics tile the retinal space with no spatial overlap or voids, except at the sparse positions where S–cones are located.

#### 2.2.2 Mosaics synthesized under different spatial compactness/spectral purity tradeoffs (stage 2B)

As mentioned previously, stage 2B allows for different optimizations of cone pooling within the mRGC RF centers, which is controlled by the spatial compactness/spectral purity tradeoff parameter, *ϕ*. Fig. 4 depicts examples of mRGC RF center mosaics all synthesized at a single eccentricity (12°), but under different values of *ϕ*. The mosaic synthesized under *ϕ* = 1, where spatial compactness is maximal and spectral purity constraint is not enforced, is depicted in Fig. 4A. Note that the RF centers tile the visual field relatively uniformly with no overlap. Figsures 4B and 4C depict mosaics synthesized as *ϕ* decreases to 0.5 and 0.0, respectively, which increasingly enforces center connections to cones of the same type. Note that this occurs at the cost of spatial compactness with the RF centers becoming spatially disordered and overlapping to a greater extent.

**Fig. 4.**
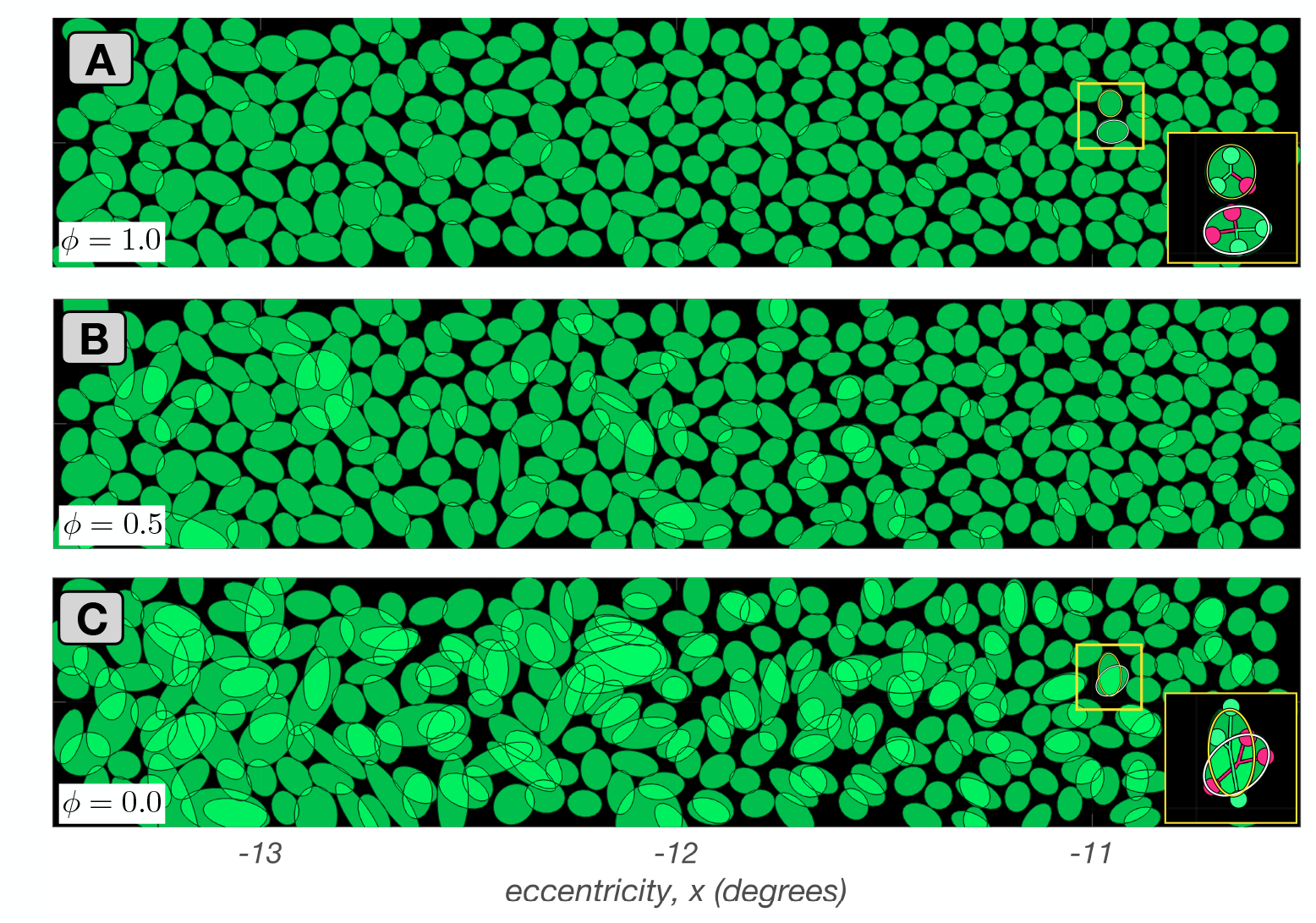
Mosaics of mRGC RF centers at the end of stage 2B. Depicted here are 3.0^*o*^ × 0.5^*o*^ mRGC mosaics, each centered at 12^*o*^ along the temporal horizontal meridian, but synthesized under different values of tradeoff between spatial compactness and spectral purity, *ϕ*. **A:** *ϕ* = 1.0 (maximal spatial compactness). **B:** *ϕ* = 0.5. **C:** *ϕ* = 0 (maximal spectral purity). Insets in A and C depict pooling of cones within the RF centers of the two mRGC RF centers contained within the yellow square. The inset in C illustrates how RF center overlap and spatial jitter is introduced as the algorithm avoids cones that are close to the RF center in order to maximize spectral purity within RF centers.

By varying *ϕ* we can examine the effect that cone-selective pooling may have on mRGC RF spatial structure, as well as on the spatio-chromatic processing in the mRGC pathway. Current electrophysiological evidence favors little selective cone pooling within peripheral RF centers of mRGCs, i.e., a *ϕ* value of ≈1 [15, 16, 33]. However, the degree of cone type selectivity in more central locations is not known with as much certainty. For example, there is anatomical evidence that ON-center mRGCs in the fovea contact multiple ON-cone bipolars, as opposed to OFF-center mRGCs, which contact single OFF-cone bipolars [34], and also electrophysiological evidence that the RFs of parafoveal mRGCs appear to be pooling from more than one cones [35].

The question of whether foveal mRGCs that pool from more than one cone in the RF centers are doing so selectively remains unanswered. Our modeling approach allows exploration of the benefits and tradeoffs of cone-selective pooling at any retinal eccentricity, although we do not pursue such exploration in this paper.

#### 2.2.3 Mosaics with divergent cone connections (stage 2C)

In the final sub-stage of establishing the mRGC RF center wiring to the cone mosaic, the mutual exclusivity constraint is lifted and single cones are permitted to connect to multiple nearby mRGC RF centers. This divergence of cone connections is enabled by replacing the binary distribution of cone pooling weights in the mRGC RF centers with a supra Gaussian distribution, as illustrated in Fig. 5. Fig. 5A depicts how a progressively increasing overlap in neighboring mRGC RF centers with eccentricity, is accomplished by varying the exponent of the supra-Gaussian distribution. In central retina, the exponent is kept at 10, which results in a flat top distribution of weights with minimal overlap between neighboring RF centers (gray histograms in the inset of Fig. 5A). As eccentricity increases beyond 7°, the exponent decreases, reaching a value of 2 (Gaussian) at around 15°, which results in Gaussian distributions of weights with significant overlap between neighboring RF centers (red histograms in the inset of Fig. 5A).

**Fig. 5.**
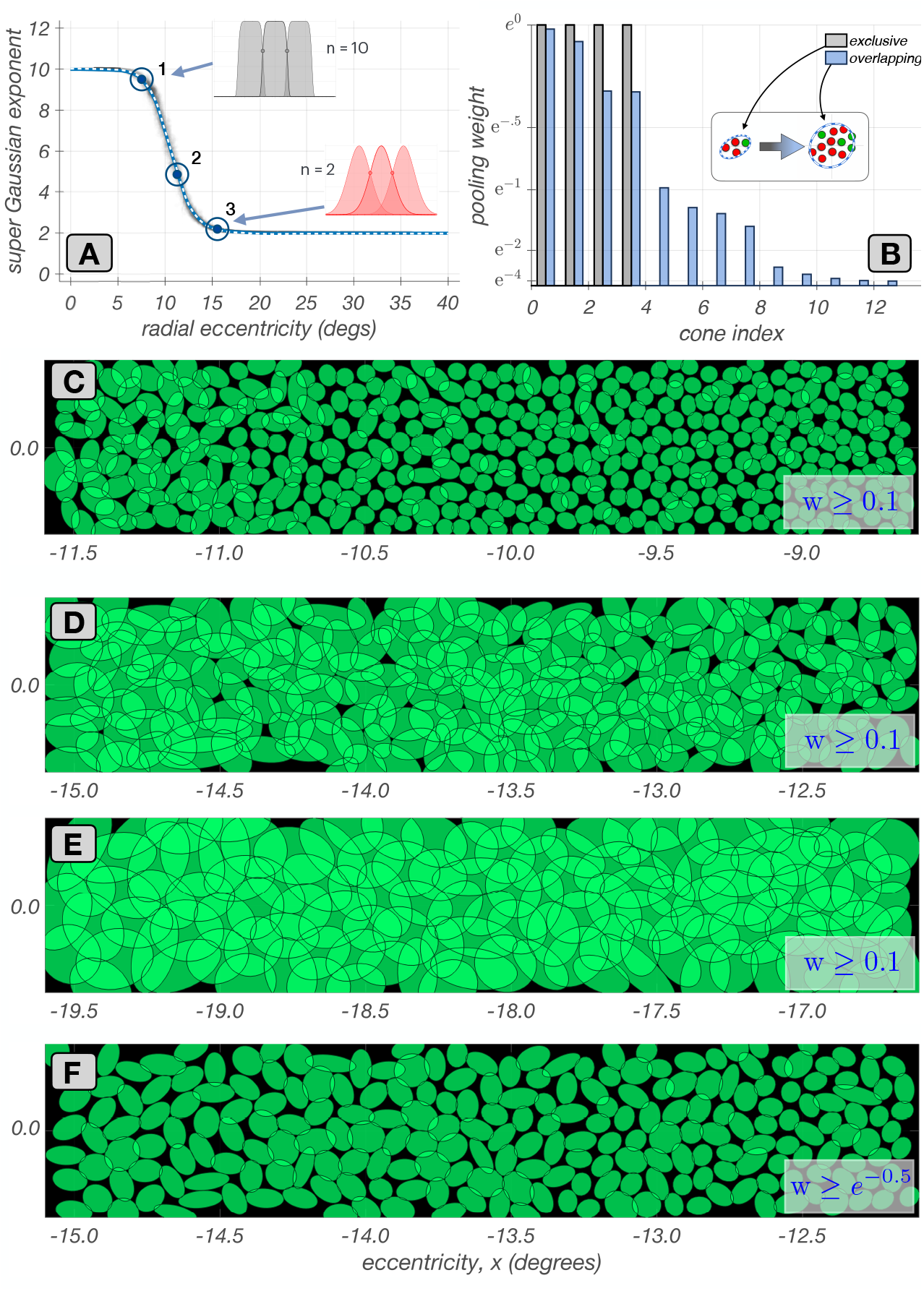
Mosaics of mRGC RF centers with divergent cone connections (stage 2C). **A:** Variation of supra Gaussian exponent with eccentricity. The exponent is set to 10 in the central retina, resulting in flat top weight distribution with zero overlap (gray histograms). As eccentricity is increased, the exponent is gradually decreased, achieving a value of 2.0, at around 15° (red histograms). **B:** Transformation of cone pooling weights, from binary, in mutually exclusive connections, (gray histogram) to non-binary in shared cone connections, (blue histogram) due to the supra-Gaussian distribution for an example mRGC. Insets depict the spatial arrangement of cones that are connected with binary and non-binary weights. **C, D, & E:** Mosaics at 10°, 13°, and 18°, along the temporal horizontal meridian with divergent cone connections. The RF center ellipses encompass the ensemble of cones with pooling weights *≥* 0.1. **F:** Same mosaic as **C**, but with ellipses showing cones with pooling weights *≥ e*^−0.5^.

This mechanism simulates the fact that in the fovea, input to mRGC RF centers comes exclusively or mostly [34, 35] from a single cone. In the periphery, *in vitro* measurements reveal that neighboring mRGC RF centers abut at approximately one standard deviation of their Gaussian RF profile [32].

The transformation of cone pooling weights from binary and mutually exclusive to graduated and shared is depicted in Fig. 5B for an mRGC located at an eccentricity of 12°, with gray and blue histograms depicting spatial distributions of cone pooling weights in this cell before and after sub-stage 2C.

Figs 5C–E depict mosaics with divergent connections synthesized at three eccentricities. In these mosaic depictions, each green ellipse represents the spatial extent that encompasses all cones that are pooled by the RF center of a single mRGC with weights ≥ 0.1. For the mosaic centered at 10° (panel C), divergence of cone connections has just begun. The overlap in RF centers due to the divergence of connections increases as we move in eccentricity from 9° on the right side to 11°, on the left side. For the mosaic centered at around 13° (panel D), cone divergence and RF center overlap is higher and again increases with increasing eccentricity. For the mosaic centered at around 18° (panel D), divergence of cone connections has assymptoted, and we have a constant RF center overlap.

Finally, Fig. 5F provides a visualization comparable to the visualization commonly reported by *in vitro* RF mapping studies [32]. It depicts the same mosaic as Fig. 5C, but with ellipses encompassing cones that are pooled with weights ≥ *e*^−1*/*2^ ≈ 0.67.

This depiction choice makes the overlap less visually salient.

### 2.3 Connecting cones to mRGC RF surrounds (Stage 3) Overview

In the last stage of mRGC mosaic synthesis, we derive the cone pooling weights for the mRGC RF surrounds. Since there are no clear anatomical data on surround sizes, these weights are determined using *in vivo* characterizations of the visual space–referred spatial transfer function (vSTF) of mRGCs in the macaque [17], which are available as a function of eccentricity.

We incorporate these data into the model using numerical optimization. More specifically, we determine the cone-to-mRGC RF surround connections such that a forward simulation of the *in vivo* physiological experiments of of Croner & Kaplan through the model best reproduces the experimental data. This approach allows us to use data collected through physiological optics, which blur the stimulus in an eccentricity and wavelength dependent manner, to determine the wiring of cones to mRGC RF surrounds.

Importantly, the optimization is achieved while adhering to the already established connectivity between the cone mosaic and mRGC RF centers. Simultaneously, the parametric form of the surrounds is constrained based on Packer & Dacey’s characterizations of the spatial RF of macaque H1 horizontal cells [36], which are the main components of the linear spatial mRGC RF surrounds [37]. The use of optimization around forward simulation of an experiment to integrate data from multiple non-commensurate sources is an important innovation of our RGC modeling approach. Stage 3 proceeds in three sub-stages.

- **Stage 3A: computation of visual space–referred cone mosaic responses to stimuli used to measure vSTFs in macaque mRGCs**. We begin by computing cone mosaic responses to gratings of different spatial frequencies delivered to the retina via human physiological optics [38], as a proxy of how macaque optics blur the stimuli. This step is necessary for us to use the *in vivo* data of Croner & Kaplan [17], which were collected with stimuli viewed through the animal’s natural optics.
- **Stage 3B: optimization of surround cone pooling functions for a subset of target synthetic mRGCs**. We choose a set of target cells that span the extent of the synthesized mRGC mosaic, and optimize the surround cone pooling functions for each. This optimization is done so that the ensuing target cells have (a) vSTF characteristics that are well approximated by a Difference of Gaussians (DoG) model, (b) have fitted DoG model parameters reasonably matching those reported by Croner & Kaplan for eccentricities corresponding to the eccentricity of the target cells, and (c) have surround cone pooling weights that maintain macaque H1-like spatial properties as characterized by Packer & Dacey.
- **Stage 3C: computation of surround cone pooling weights for all cells in the synthesized mRGC mosaic**. Once the surround cone pooling functions are derived for the target synthetic mRGCs, surround pooling weights for all cells in the synthesized mRGC mosaic are computed by interpolation. A small amount of jitter in the ratio of the surround to center weights is added to simulate the variance in integrated surround to center ratio at each eccentricity seen in the macaque data.

#### 2.3.1 Computation of visual space–referred cone mosaic responses to stimuli used to measure vSTFs in macaque mRGCs (Stage 3A)

We employ the ISETBio machinery to compute the excitation of the input cone mosaic to achromatic gratings of different spatial frequencies delivered to the retina via physiological optics. This process captures several crucial spatio-chromatic effects in the transformation of scene radiance into cone responses: spatial and chromatic filtering by physiological optics, spectral filtering by the eye’s inert pigments, and sampling by the interdigitated trichromatic cone mosaic. To mimic the phototransduction process, cone excitation responses are converted to cone modulation responses.

In these computations, we employ human physiological optics matched to the eccentricity of each synthesized mRGC, but we adjust the defocus term of the modeled optics so as to maximize the Strehl ratio (ratio of peak sensitivity of the PSF at the wavelength of focus, here 550 nm, to the peak sensitivity of a diffraction-limited PSF). This is done as a proxy to the experimental paradigm of Croner & Kaplan, in which corrective lenses were used to maximize cell responses at high spatial frequencies (personal communication with the late Ehud Kaplan).

#### 2.3.2 Deriving surround cone pooling functions for a subset of target synthetic mRGCs (Stage 3B)

Croner & Kaplan reported summaries of spatial RF characteristics across populations of mRGCs at different eccentricities, which were derived by fitting a Difference of Gaussian (DoG) model to the measured vSTFs, and tabulated the variation of the DoG parameters with eccentricity. The DoG model defined in spatial frequency, *ω*, domain is given by:

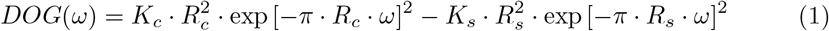

where *K*_*c*_ and *K*_*s*_ are the peak sensitivities of the RF center and RF surround mechanisms, and *R*_*c*_ and *R*_*s*_ are the corresponding characteristic radii.

The vSTF of a typical macaque mRGC is depicted in Fig. 6A with cyan disks. The solid heavy line depicts the fitted DoG model, with the component center and surround vSTFs depicted by the thin solid and dashed lines, respectively. The shape of the vSTF is determined by two key measures, the ratio of surround to center characteristic radii, *R*_*s*_*/R*_*c*_, and the ratio of surround to center integrated sensitivities, *K*_*s*_*/K*_*c*_ × (*R*_*s*_*/R*_*c*_)^2^. The distributions of these two ratios as a function of eccentricity in the population of mRGCs recorded by Croner & Kaplan are depicted by the gray squares in Figs 6B1 & 6B2. The mean variation in these two ratios, shown as dashed lines, are the target values used to derive the surround cone pooling weights in the synthetic mRGCs.

**Fig. 6.**
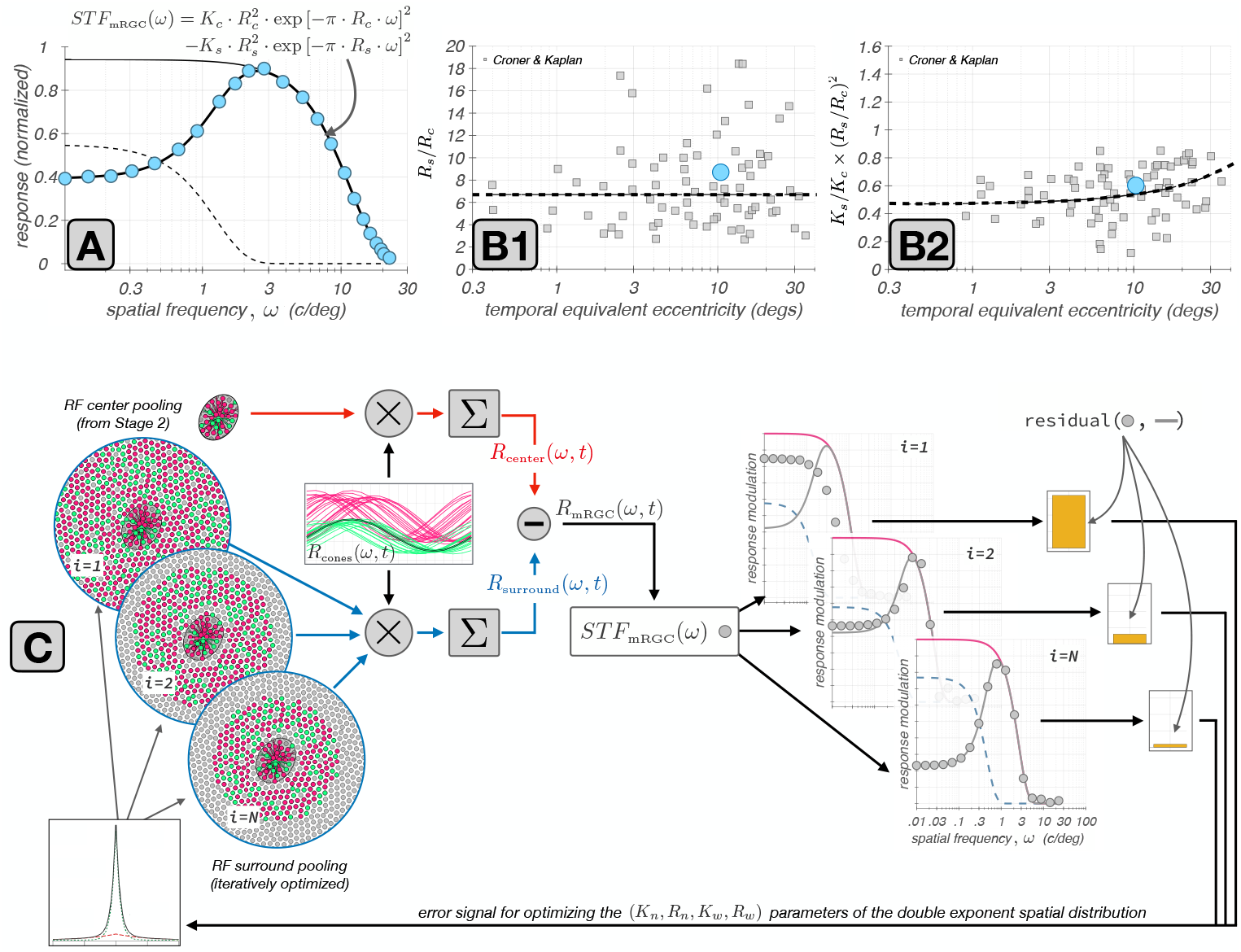
Derivation of cone weights to mRGC surrounds. **A:** Typical macaque mRGC vSTF (cyan disks) fitted with a Difference of Gaussians (DoG) model (thick black line). The component center and surround vSTFs are depicted by the thin black and the dashed lines, respectively. **B1 & B2:** Ratios of surround to center characteristic radii, *R*^*s*^*/Rc*, and ratios of surround to center integrated sensitivities, 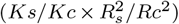 as a function of eccentricity in the population of mRGCs recorded by Croner & Kaplan [17]. The dashed lines represent the trends in these two ratios as a function of eccentricity. The cyan disks depict the ratios for the example vSTF depicted in A. **C:** Depiction of the iterative estimation of surround cone pooling weights by forward simulation of the Croner & Kaplan vSTF measurements. See description in text.

The optimization process is illustrated schematically in Fig. 6C. The vSTF of the target synthetic mRGC is computed by forward simulation of the experiment of Croner & Kaplan. The time courses of responses of L– and M–cones from the input cone mosaic to a drifting grating stimulus of spatial frequency *ω, R*_cones_ (*ω, t*), (computed in Stage 3A) are depicted by the red and green traces in the rectangular panel of Fig. 6C. A spatially weighted sum of these cone responses using the RF center cone pooling weights (computed in Stage 2), is used to compute the response of the RF center, *R*_center_(*ω, t*). This operation, which is depicted by the red computation arm in Fig. 6C, is fixed throughout the optimization of the surround.

In the computation of the spatial distribution of surround cone pooling weights, we impose a parametric form that is described by the sum of a narrow and a wide exponential components, based on characterizations of the spatial RF properties of H1 horizontal cells by Packer & Dacey [36]:

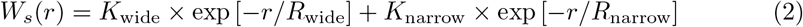

where *r* is the radial distance from the RF center, *K*_*wide*_ and *K*_*narrow*_ are the peak sensitivities of the wide and the narrow components, respectively, and *R*_*wide*_ and *R*_*narrow*_ are the corresponding characteristic radii.

Beginning with a random initial value for the parameters of the double exponential distribution, we compute an initial estimate of the surround cone weights by evaluating *W*_*s*_(*r*) at the vicinity of the input cone mosaic that surrounds the RF center. These weights are depicted in the top-left circular panel of Fig. 6C (labeled as *i* = 1, with *i* denoting iteration). Using these initial weights we compute a weighted sum of the surround cone responses to derive the initial estimate of the surround response, *R*_surround_(*ω, t*) This operation is depicted by the blue computation arm in Fig. 6C.

The composite response of the synthesized mRGC is obtained by instantaneously subtracting the surround response from the center response, as follows:

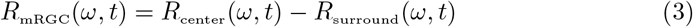

The amplitude modulation of *R*_mRGC_(*ω, t*) is taken as the value of the vSTF at the examined spatial frequency, *STF*_mRGC_(*ω*). Repeating over a range of spatial frequencies, we obtain the initial estimate of the full vSTF, which is depicted by the gray disks in the top-right rectangular panel of Fig. 6C, labeled as *i* = 1.

Following the experimental procedure of Croner & Kaplan, we fit the computed *STF*_mRGC_(*ω*) with a DoG model. This fit is depicted by the solid gray line in the top-right rectangular panel of Fig. 6C. Note that the DoG model fit is constrained so that its shape parameters, *R*_*s*_*/Rc*, and 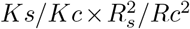, both lie within a narrow range of the mean values of *R*_*s*_*/Rc*, and 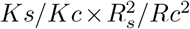 ratios reported for macaque mRGCs at corresponding eccentricities [17]. At the first iteration, the residual between the computed vSTF and the DoG model fit to it, is large.

This residual, depicted by the yellow bar in the right-most panel of Fig. 6C, serves as an error signal. The optimization algorithm minimizes this error signal by adjusting the parameters of *W*_*s*_(*r*), which controls the surround weights, with the additional constraint that the parameters of *W*_*s*_(*r*) remain within a range of the values reported in macaque H1 horizontal cells [36].

The optimized surround cone pooling function is obtained when the residual between the computed and fitted vSTFs reaches a minimum value, labeled as *i* = *N*, in Fig. 6C. Additional details about this surround optimization method is provided in Section B.1.

### 2.4 Deriving surround cone pooling weights for each cell in the mosaic(stage 3C)

The optimization of the surround cone pooling functions is a computationaly expensive process. It is therefore conducted on a sparse spatial grid (with *N*_*xy*_ nodes), which encompasses the spatial extent of the synthesized mRGC mosaic. At each node of the spatial grid, we determine the range of cone numerosities in the RF centers of nearby synthetic mRGCs, and we derive optimized surround cone pooling functions for each of the encountered RF center cone numerosities (*N*_*c*_), and we do this twice, once for L-cone dominated RF centers, and once for M-cone dominated RF centers.

Once these *N*_*xy*_ *× N*_*c*_ *×* 2 surround cone pooling functions are derived, we compute surround cone pooling weights for all synthetic mRGCs. For each target mRGC we determine the 3 nearest spatial grid nodes, and extract the optimized surround cone functions that were derived at this node for the cone numerosity that matches that of the target mRGC, for both L- and M-center cone dominance variants. Then we evaluate the six optimized surround pooling functions at the input cone mosaic in the vicinity of the target mRGC, deriving six sets of surround cone pooling weights. The target cell’s surround cone pooling weights are determined by interpolating the 6 sets of weights spatially, weighted inversely proportionally by the distance between the location of the target mRGC and the location of the optimized model, and spectrally, weighted based on the relative L-/M-cone weight ratio in the RF center of the target mRGC.

#### 2.4.1 Adjusting the surround pooling variance

The final step in the generation of the mRGC RF surrounds is to apply a noisy scalar multiplier to all surround pooling weights of individual mRGCs. The value of this scalar is chosen so that the variance in the ratio of surround to center integrated sensitivities, *K*_*s*_*/K*_*c*_ *×*(*R*_*s*_*/R*_*c*_)^2^, of the synthetic mRGCs matches the variance observed in the population of macaque mRGCs recorded by Croner & Kaplan. The manipulation in *K*_*s*_*/K*_*c*_ *×* (*R*_*s*_*/R*_*c*_)^2^ variance does not require re-computing the surround pooling functions. This is unlike manipulating the variance in the *R*_*s*_*/R*_*c*_ ratio, which requites re-computing the surround pooling functions.

#### Computing mRGC responses from cone mosaic responses

A fully synthesized mRGC mosaic consists of two connectivity matrices: **W**_center_(*i, k*), determined in synthesis stage 2, and **W**_surround_(*i, k*), determined in synthesis stage 3, which capture the weights by which the RF center and surround, respectively, of the *k*^*th*^– cell in the mRGC mosaic pools signals from the *i*^*th*^ cone in the input cone mosaic. Since the current version of the mRGC model does not contain a temporal component, the response of the *k*^*th*^– mRGC to some stimulus at time instant, *t, R*_stim_(*k, t*), is computed instantaneously from the response of the input cone mosaic to that stimulus at time *t*, **C**_stim_(:, *t*), as follows:

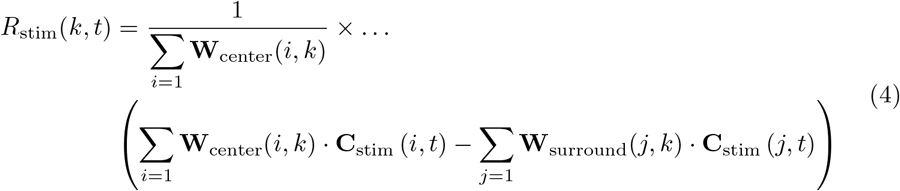

To mimic adaptation to the background stimulus, the mRGC mosaic typically operates on cone contrast responses, instead of cone excitation responses, so the **C**_stim_(:, *t*) term in the above equation is computed as follows:

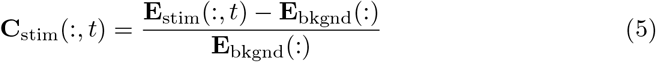

where **E**_stim_(:, *t*) is the cone mosaic excitation response to the examined stimulus at time *t*, and **E**_bkgnd_(:) is the cone mosaic excitation response to a uniform field, zero contrast stimulus, whose mean (*x, y*) chromaticity and mean luminance match those of the examined stimulus.

## 3 Results

key feature of our model is its dual representation of mRGC receptive field (RF) properties, which separates neural circuitry from optical effects. The first representation, in *retinal space*, models the direct pooling of cone signals by the RF center and surround. This describes the cell’s intrinsic spatio-chromatic filtering and is directly comparable to anatomical data and physiological measurements that bypass the eye’s optics (e.g., *in vitro* or adaptive optics experiments [10, 39]). In contrast, the second representation, in *visual space*, models the end-to-end processing of a stimulus as it passes through the eye’s optics to the mRGC mosaic. This representation is applicable to conventional *in vivo* physiology and psychophysical assessments of visual function.

The ability to go back and forth between cone and visual space is critical to understanding how retinal cone pooling interacts with physiological optics to generate the processing characteristics of cells in visual space, which is what ultimately determines natural visual performance. This ability is also critical in interpreting results from *in vivo* physiology in terms of the underlying retinal wiring [40], as well as to relating results obtained under adaptive optics viewing conditions to results obtained under natural viewing conditions [10].

In this section we illustrate and contrast spatial RF characteristics of synthetic mRGCs in the two representations and validate the properties of synthetic mRGCs against those of macaque mRGCs as characterized by *in vivo* and *in vitro* physiological studies.

### 3.1 Spatial characteristics of synthesized mRGC receptive fields

Spatial characteristics of cells in an mRGC mosaic synthesized at 4.5^*o*^ along the temporal horizontal meridian are depicted in Fig. 7. The employed mosaic is depicted in Fig. 7A. The numbered positions in Fig. 7A identify the locations of three selected cells whose spatial RF characteristics are explored in detail next. The cone pooling maps of these exemplar mRGCs are depicted in Figs 7B1–B3. Here, pink and cyan disks depict L– and M–cones, respectively, that are pooled by the RF center with a weight ≥ 0.1, or by the RF surround with a pooling weight ≥ 0.005, and gray disks depict cones that are either not pooled at all or pooled with a weight less than the threshold for labeling. The solid and dashed lines depict the spatial pooling extents of the RF center and surround mechanisms, respectively.

**Fig. 7.**
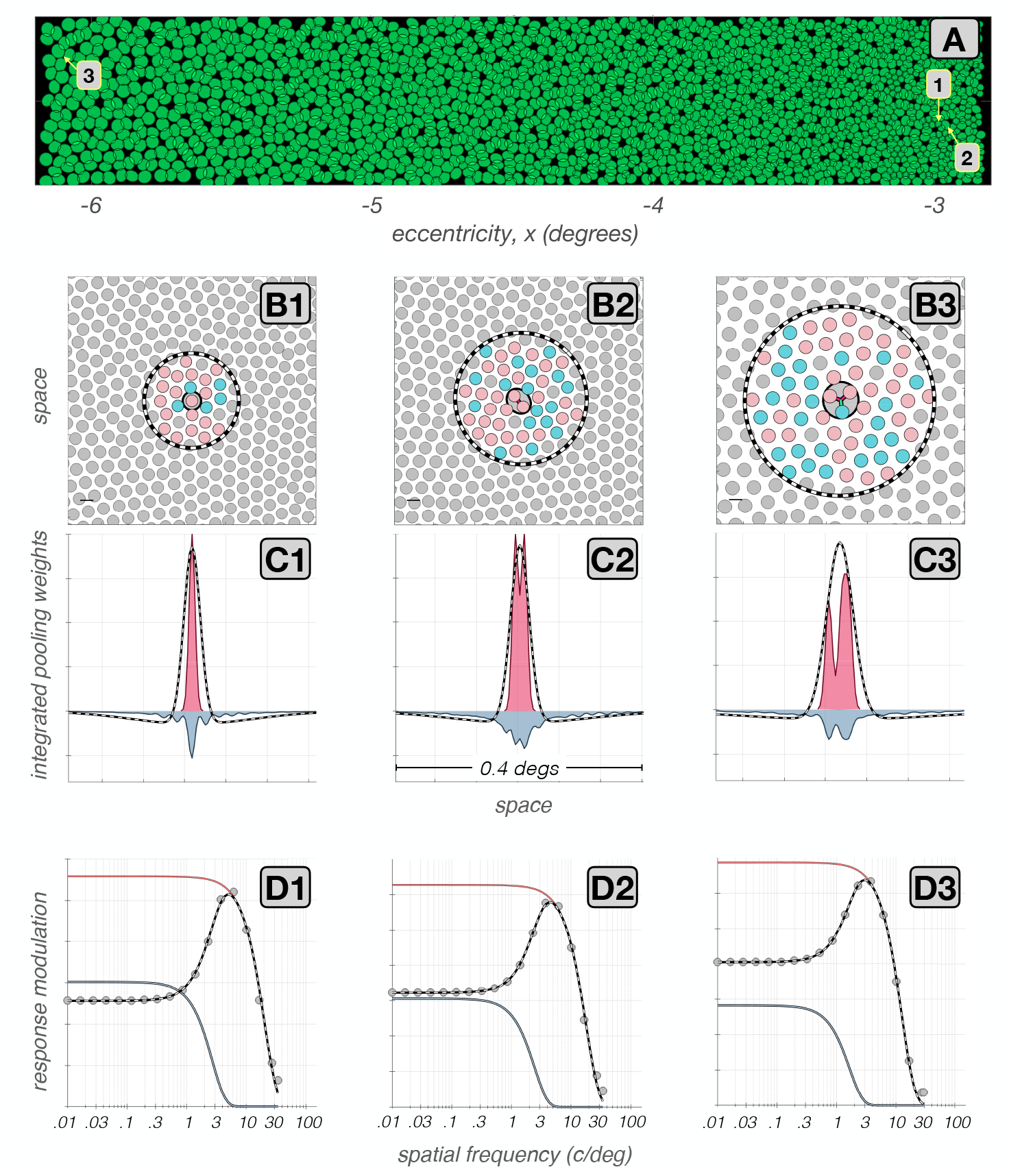
Spatial RF characteristics of synthetic mRGCs. **A:** Mosaic of RF centers of an mRGC mosaic synthesized at 4.5° along the temporal horizontal meridian. **B1–B3:** Cone pooling maps for 3 exemplar cells whose positions within the mRGC mosaic are labeled in A. Pink and cyan disks depict L– and M–cones, respectively, whose RF center pooling weights are ≥ 0.1, or whose RF surround weights are ≥ 0.005. Gray disks represent either S-cones, which are not pooled in our model, or L–/M–cones with pooling weights lower than the labeling thresholds. The solid and dashed black lines depict the extents of the RF center and surround pooling regions. **C1–C3:** Y-axis integrated cone pooling weight profiles within the RF center (maroon) and surround (slate). The dashed lines depict the visual space–referred line weighting functions as derived by fitting Difference of Gaussians (DoG) models to each cell’s vSTF. **D1–D3:** The vSTFs of the exemplar mRGCs, computed under physiological optics, are depicted by the gray disks. The gray, maroon, and slate lines depict the DoG model fits to these vSTFs, and the models’ component center and surround vSTFs, respectively.

The cell depicted in Fig. 7B1 is located at an eccentricity of 3^*o*^. Its RF center pools from a single L-cone and its RF surround pools from a total of 16 L– and M– cones. The cell depicted in Fig. 7B2, also located at 3^*o*^, pools from two L–cones in its RF center, and its RF surround is larger, pooling from 44 L– and M–cones. The cell depicted in Fig. 7B3 is located at 6^*o*^. Its RF center, which pools from 2 L–cones and 1 M–cone, and its surround are both larger than those of the first 2 cells.

The cone pooling maps depicted in Figs 7B1–B3 illustrate the spatial connectivity between the input cone mosaic and the center and surround subregions of mRGC RFs, but do not depict the strength of these connections. In this sense, these maps depict the type of information that is available from detailed anatomical studies.

Figs 7C1–C3 add to this view by providing information about the strength of the cone inputs for the three exemplar cells. Here, the maroon and slate histograms depict the cells’ spatially integrated (along the y-axis) cone pooling weights for the RF center and the surround mechanisms, respectively. Note that in the cell depicted in Fig. 7C1, the double exponential spatial profile of the surround cone pooling mechanism, with a sharp peak around the RF center and more shallow weights in peripheral regions, is prominent. In the two other cells shown, this feature is less prominent.

This observation, where cells with larger RF centers have less peaked surround weights than cells with smaller RF centers is seen commonly in our synthetic mRGCs. The change in surround pooling characteristics results from constraints in the model, which maintain vSTF shape parameters that are consistent *in vivo* measurements [17] while at the same time remaining consistent with the surround parametric form indicated by measurements of H1 receptive fields [36].

Visual space–referred spatial transfer functions (vSTFs) are commonly measured in *in vivo* physiological assessments to estimate spatial RF properties of mRGCs [17, 18]. The vSTFs computed for the three examined synthetic mRGCs are depicted by the gray disks in Figs 7D1–7D3. The corresponding DoG model fits are depicted by the solid gray lines, and the spatial RF profiles corresponding to these DoG model fits are depicted by the dashed lines in Figs 7C1–C3. Contrasting these inferred spatial RF profiles with the actual cone pooling profiles, it becomes evident that one cannot use characterizations obtained under physiological optics viewing to directly infer the characteristics of spatial pooling of cone signals in the retina. We discuss this issue further in later sections.

### 3.2 Validation against *in vivo* physiology across the visual field

To validate our model, we synthesized mRGC mosaics across a wide region of the retina, and computed vSTFs of individual mRGCs by probing them with drifting achromatic gratings of different spatial frequencies delivered to the retina under human physiological optics appropriate for the eccentricity of the examined cells, simulating the experimental paradigm of Croner & Kaplan [17]. To compare synthetic and macaque mRGCs we fitted the synthetic cell vSTFs with the DoG model employed by Croner & Kaplan and compared the ratios of surround to center characteristic radii, *R*_*s*_*/Rc*, and ratios of surround to center integrated sensitivities, 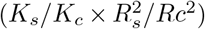, to those reported by Croner & Kaplan [17]. The results of this analysis are depicted in the left and right panels of Fig. 8 for mRGC mosaics synthesized under the physiological optics of two different human observers. Figs 8A1 and 8A2 depict the comparison in the distribution of *R*_*s*_*/Rc* ratios. Gray squares depict the macaque mRGC data and the blue density plots depict the 5%–95% percentile range of corresponding parameters in a population of 66,128 synthetic mRGCs. The three yellow disks in Fig. 8A1 correspond to the three exemplar cells illustrated in Fig. 7. Note that the *R*_*s*_*/Rc* ratios in synthetic mRGCs follow the macaque data across eccentricity for both human subjects. The synthetic data do not, however, capture the full variance seen in the macaque data, as is evident by the marginal histograms (Fig. 8A3). To capture the full variance seen in the macaque *R*_*s*_*/Rc* ratios, we could consider synthesizing multiple surround pooling functions, each with different target values of *R*_*s*_*/R*_*c*_, and then randomly selecting for each synthesized mRGC from the multiple sets.

**Fig. 8.**
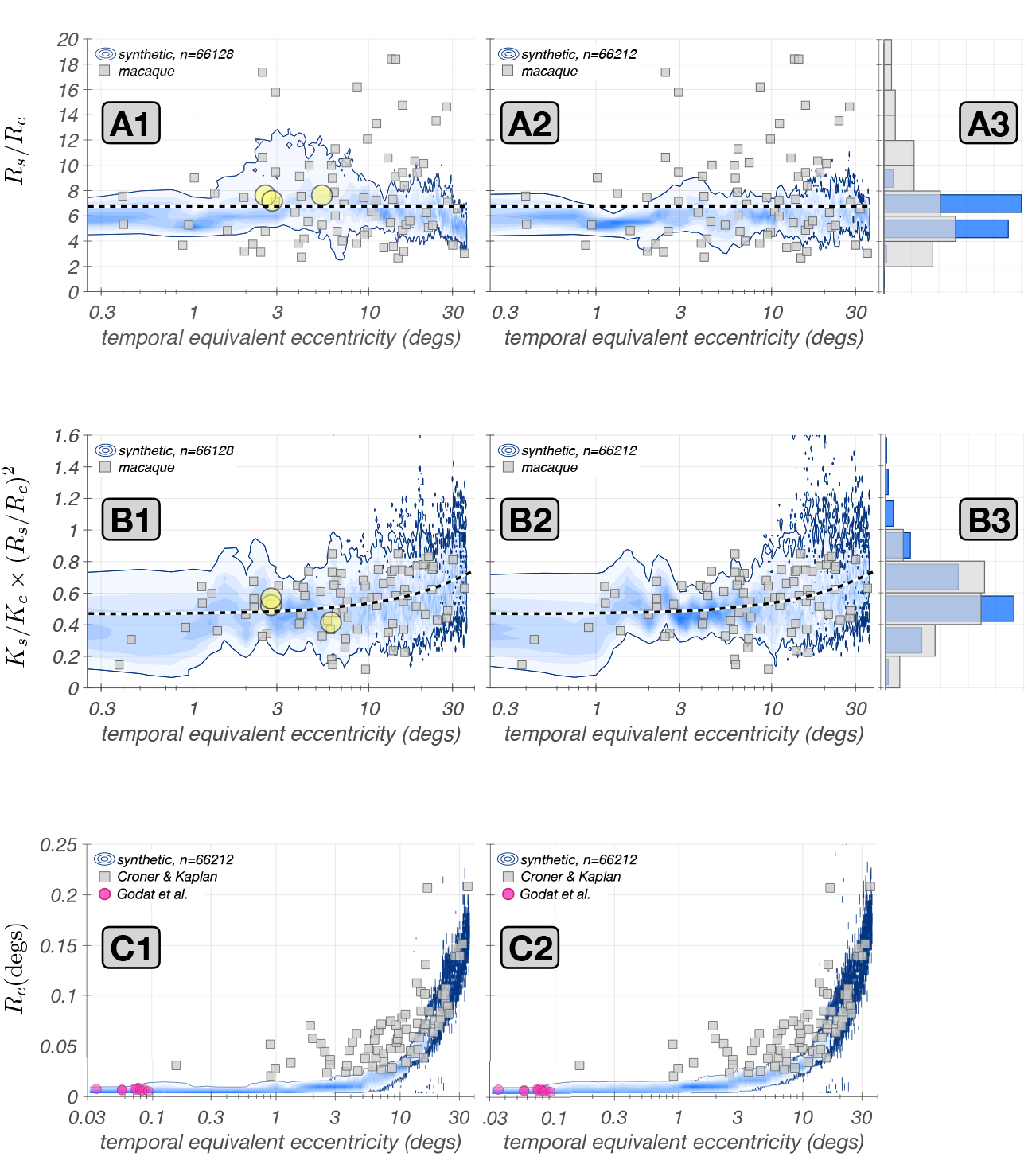
Validation against *in vivo* measurements. In all panels, gray squares depict data from the population of macaque mRGCs recorded by Croner & Kaplan [17]. Blue contours depict the probability density function of the examined parameter in a population of 66,128 synthetic mRGCs with color saturation encoding probability level. Solid blue lines represent the 5% – 95% percentile range of examined parameter. Left and right panels are for mosaics synthesized under physiological optics of two different human subjects. **A1–A2:** Correspondence in ratio of surround–to–center characteristic radii, *R*_*s*_*/R*_*c*_, across eccentricity. The dashed line represents the target value that is in effect during the optimization of the synthetic mRGC surrounds, which is the mean value of *R*_*s*_*/R*_*c*_ across the population of all mRGCs recorded by Croner & Kaplan. **A3:** Marginal histograms of *R*_*s*_*/R*_*c*_ for macaque (gray) and synthetic mRGCs (blue). **B1–B2:** Correspondence in ratio of surround–to– center integrated sensitivities, *K*_*s*_*/K*_*c*_ *×* (*R*_*s*_*/R*_*c*_)^2^, across eccentricity. The dashed line represents the target values in effect during the optimization of the synthetic mRGC surrounds, which is the trend observed with eccentricity in the population of the macaque mRGCs recorded by Croner & Kaplan. **B3:** Marginal histograms of *K*_*s*_*/K*_*c*_ *×* (*R*_*s*_*/R*_*c*_)^2^ for macaque (gray) and synthetic mRGCs (blue). **C1–C2:** Correspondence in RF center characteristic radius, *R*_*c*_, across eccentricity. The fuschia disks represent the *R*_*c*_ of foveolar mRGCs recorded by Godat *et al*. [10], back-projected in visual space using the monkey’s own physiological optics.

On the other hand, the integrated sensitivity rations, 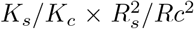, of the synthetic mRGC population, depicted in Figs 8B1–B3, capture both the trend with eccentricity and the variance of the macaque data. The variance match was achieved by enforcing a target variance in the 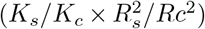 ratio of the synthetic cells as described earlier.

Note that, although we did use the mean variation with eccentricity of macaque *R*_*s*_*/R*_*c*_ and 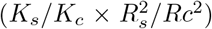 ratios during construction of the model, the model was derived using additional constraints: those imposed by the densities of cones and mRGC RFs, by the spatial characteristics of H1 horizontal cells, and by measurements of human optics. These validations, therefore, check both that we have not over constrained our model in a manner that makes it inconsistent with the macaque data, and that our method of interpolating surround pooling weights from models derived at a set of discrete retinal locations works well.

We next examined the correspondence between synthetic and macaque mRGCs in their visual space–referred RF center sizes, *R*_*c*_. Recall that in synthesizing mRGC mosaics, the RF centers are constructed independently of the Croner & Kaplan physiological data, using only anatomical data and estimates of RF center overlap obtained from *in vitro* physiology in the periphery [32]. The comparison between synthetic and macaque *R*_*c*_ values is depicted in Figs. 8C1–C2. Note that the distribution of *R*_*c*_ in the synthetic mRGCs follows the trend seen in macaque mRGCs with eccentricity, with good agreement at eccentricities above 10^*o*^ for both subjects. In more central locations, however, the synthetic mRGC RF center sizes are 2–3 times smaller than those in the macaque.

We believe that the discrepancy at central locations is not a deficiency of our model, but rather results from several factors. First, the cone mosaic in our model has a peak density of 288,000 cones/mm^2^ which is near the high end of densities reported in humans [41], whereas the average macaque peak cone density is around 200,000 cones/mm^2^ [42, 43]. The higher cone mosaic density in humans implies smaller cone apertures, which in turn would bias our synthetic mRGCs towards somewhat smaller RF centers.

Second, in acute macaque experiments, optical refraction is not necessarily perfect, so there could be residual blur due to errors in refraction, as well as due to corneal edema from the contract lens used in typical multi-day acute experiments. This would increase the size of the RF centers in the physiological data relative to those in our model in central retina. Moreover, residual eye movements can occur in acute experiments, despite the ocular muscle paralysis (personal observations by N.P. Cottaris). Such residual movements would artificially enlarge estimates of RF center size for central retina mRGCs.

Finally, in the macaque experiments of Croner & Kaplan, stimulus orientation was not optimized to match any orientation bias in the RF of macaque mRGCs (Lisa Croner, personal communication), whereas in the simulated experiments, stimulus orientation was matched to the cell’s visual-space referred orientation bias, which results in the smallest possible estimate of RF center size.

Support for our assertion is provided by *in vivo* data from foveal macaque mRGC STFs obtained under adaptive optics viewing conditions [10]. The center sizes of these cells, blurred by the optics measured for the monkey subjects studied, are depicted by the purple disks in Figs. 8C1 & Figs. 8C2. Note that these align well with the *R*_*c*_ values of our synthetic mRGCs.

### 3.3 Validation against *in vitro* physiology in the periphery

We also compared spatial RF properties of synthetic mRGCs against macaque data from *in vitro* mRGC recordings. Since the *in vitro* data are not subject to optical blur, they may be compared directly to the retinal-space characteristics of our model. Data of this sort are currently only available in the peripheral retina.

The first study considered is that of Gogliettino *et al*. [44], in which the spatial RFs of mosaics of macaque mRGCs were mapped using white noise stimulation. To simulate their experiments, we probed synthetic mRGCs with white noise modulated achromatic checkerboard stimuli delivered to the retina under diffraction limited optics. To compute the spatial RFs of synthetic mRGCs, we cross-correlated the synthetic mRGC responses with the white noise stimulus sequence. Results of this analysis are depicted in Fig. 9.

**Fig. 9.**
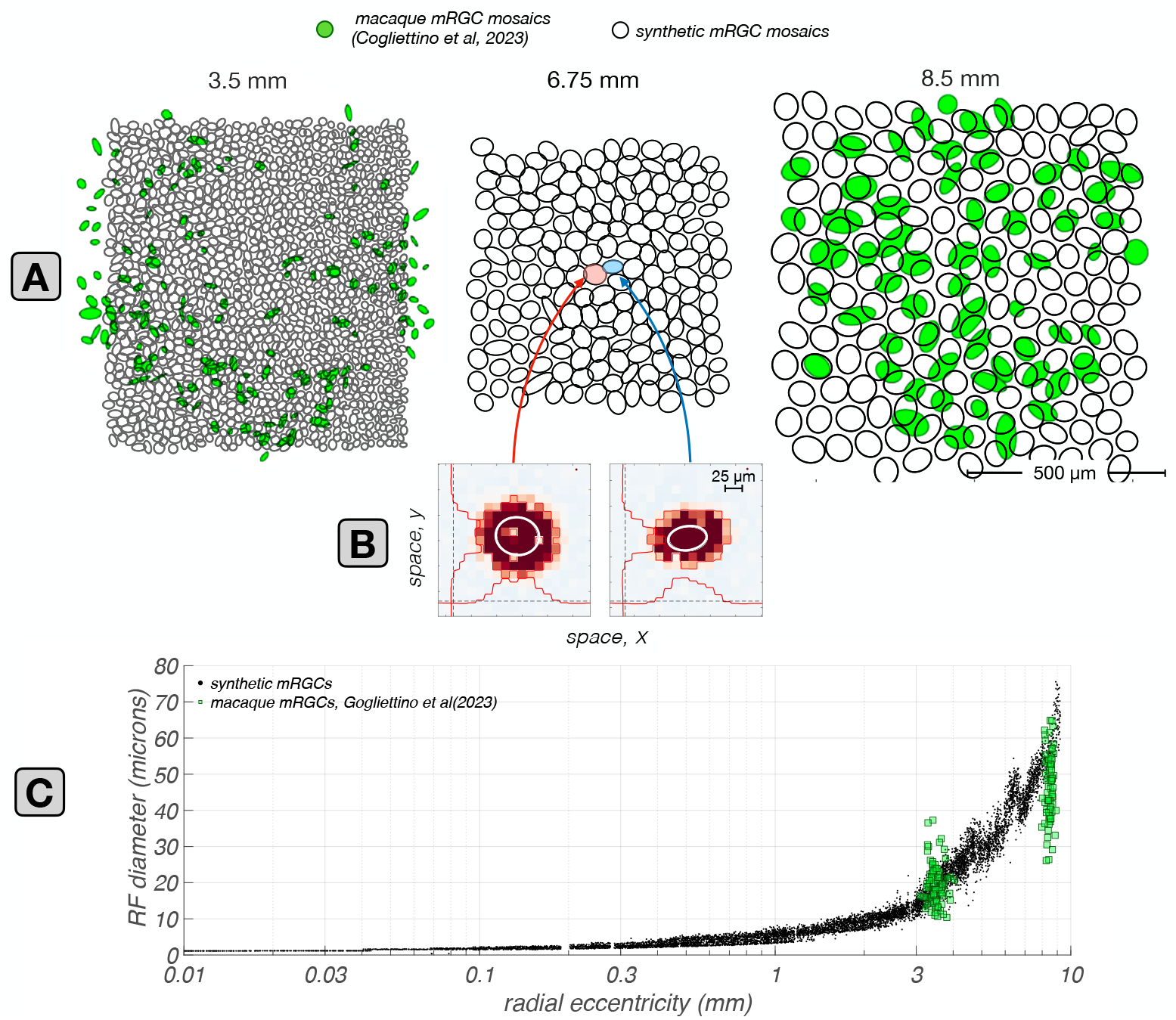
Retinal space–referred RF center sizes: synthetic vs. macaque mRGCs recorded *in vitro*. **A:** Mosaics of synthetic mRGCs synthesized at three eccentricities, 3.5, 6.75, and 8.5 mm along the temporal meridian. The black contours depict Gaussian ellipsoid fits to the increment-excitatory regions of the computed RF maps, drawn at the *e*^−1^ normalized sensitivity level. Only the increment-excitatory region of the RF map is fitted. Green contours depict RF maps from two macaque mRGCs mosaics from the *in vitro* recordings of Gogliettino *et al*. [44]. **B:** Example spatial RF maps of two synthetic mRGCs located at 6.75 mm, computed via white noise stimulation delivered to the retina under diffraction limited optics. Regions excitatory to light increments, i.e. the RF centers, and to light decrements, i.e., the RF surrounds, are indicated by red and blue colors, respectively. The scattered zero excitation spots within the light-increment RF centers correspond to the location of S-cones. White lines depict iso-contour plots of Gaussian ellipsoids fitted to the light increment–excitatory RF center region, drawn at the *e*^−1^ normalized sensitivity level. **C:** Comparison of synthetic against macaque mRGC RF center sizes across eccentricity. Black dots depict the RF diameters of synthetic mRGCs, computed from the Gaussian ellipsoid fits as 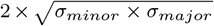, and green squares depict the RF diameters of macaque mRGCs at the two eccentricities where the *in vitro* measurements are available.

The spatial RFs of cells in synthetic mRGC mosaics at three eccentricities, 3.5 mm, 6.75 mm and 8.5 mm, are illustrated by the black ellipses in the three top panels of Fig. 9A. The superimposed green filled ellipses depict spatial RFs from macaque mRGC mosaics located at 3.5 mm and 8.5 mm. Note that at both eccentricities, there is good correspondence in RF center size and coverage between the synthetic and the macaque mRGC mosaics.

To quantify the retinal space–referred RF center sizes in synthetic mRGCs, we computed the diameter of their RF centers as 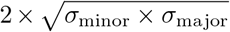, where *σ*_minor_ and *σ*_major_ are the standard deviations of the fitted Gaussian ellipsoid along its minor and major axes. The results of this analysis across eccentricity are depicted by the black dots in Fig. 9C, along with the RF center sizes of mosaics of macaque mRGCs located at 3.5 mm and 8.5 mm, which are depicted by the green squares.

Note that the correspondence between synthetic and macaque data is excellent at 3.5 mm, whereas at 8.5mm, the RF diameters of the synthetic mRGCs are, on average, 30–40% larger than the RF diameters of macaque mRGCs. The deviation in RF size at the far periphery may occur because human and macaque retinas differ somewhat in the periphery. For example, in the human retina, cone density does not change much for eccentricities *>* 5mm, whereas in the macaque retina it continues to drop as eccentricity increases [45]. The RF size deviation we observe could be the result of a higher mRGC density in the peripheral macaque retina, relative to the human retina. The second *in vitro* study we validated our synthetic mRGCs against, is that of Field *et al*. [15], which examined the spatial layout of single cone inputs to the RF centers and surrounds in peripheral macaque mRGCs. Results of this comparison are depicted in Fig. 10. Fig. 10A depicts the cone pooling maps of three synthetic mRGCs at a temporal eccentricity of 6.75 mm, and Fig. 10B depicts the spatial distribution of cone pooling weights measured by Field *et al*. [15]) in three macaque mRGCs at the same eccentricity (adapted from their Fig. 4). The visualized surround cones have pooling weights *>* 0.005 *×* the peak center cone weight for both the synthetic mRGCs and the macaque mRGCs (Greg Field, personal communication).

**Fig. 10.**
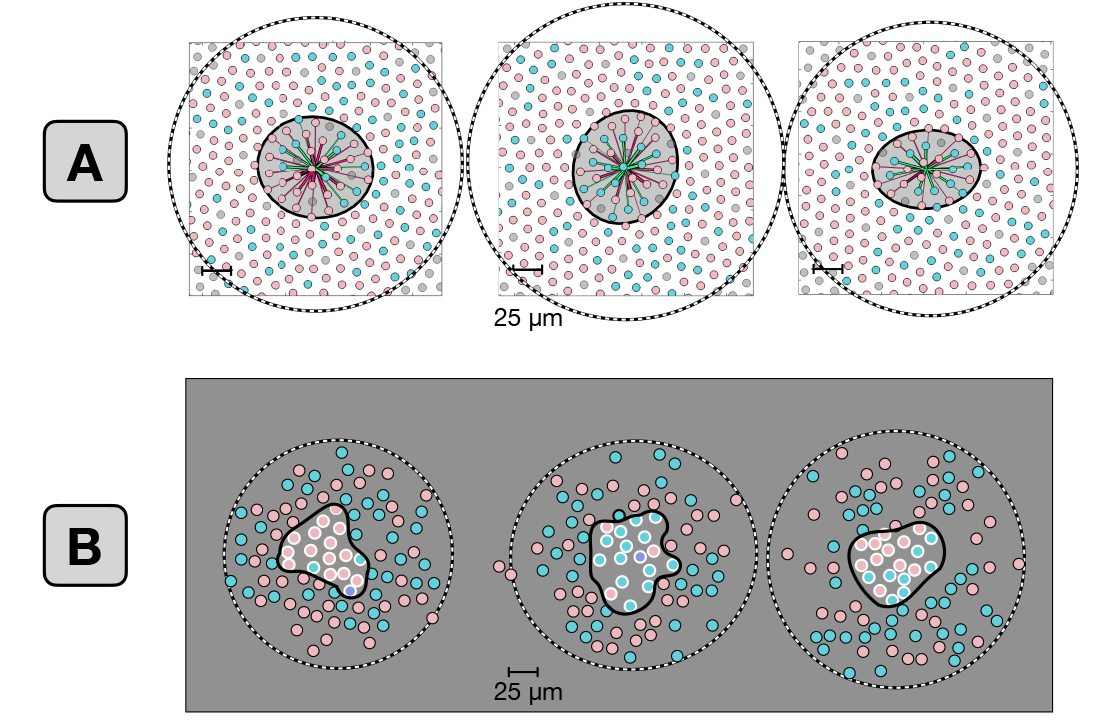
Cone pooling maps in RF centers and surrounds: synthetic vs. macaque mRGCs recorded *in vitro*. **A:** Center and surround cone pooling weight maps for three synthetic mRGCs at an eccentricity of 6.75 mm along the temporal raphe. Solid and dashed contours include cones pooled by the RF center and RF surround, respectively, with pooling weights *>* 0.005 *×* the peak center weight. **B:** Center and surround cone pooling weights for three macaque mRGCs recorded in vitro at an eccentricity of 6.75 mm along the temporal raphe. White and black disks indicate cones pooled by the RF center and RF surround respectively, with same threshold pooling weights as in A. The macaque mRGCs are from the *in vitro* recordings of Field *et al*. [15].

Note the general agreement between synthetic and macaque mRGCs in the extent of both their RF centers and surrounds, although again, synthetic mRGCs appear to have slightly larger RFs that their macaque counterparts. Also notable is that the density of cones in the synthetic mRGC cone pooling maps is higher than that seen in the macaque mRGCs. This occurs because our model is based on human cone mosaics, and human cone density is higher than macaque cone density at temporal eccentricities above 5 mm [45], which is where these comparisons are made.

This observation highlights an inherent issue in building our mRGC model, namely that we had to employ a mixture of human and macaque data sources: human data regarding the density of cones and the density of mRGC RFs across visual space, human data regarding the characteristics of physiological optics across the retina, and macaque data regarding the spatial characteristics of mRGC RFs and of H1 horizontal cells, with our validations done against macaque data. This is not ideal, as there are some differences between human and macaque retinas [45]. But, it is unavoidable given the lack of complete data in either species. The modeling framework that we devised however, which incorporates data from different sources, can be easily modified as new data become available.

### 3.4 Visual *vs*. retinal space– referred RFs: the impact of physiological optics

In this section we characterize how physiological optics interacts with the retinal cone pooling within the RFs of mRGCs to shape their visual space–referred RF properties. Fig. 11 illustrates examples of this interaction at five horizontal eccentricities, *x* = [−16^*o*^, −8^*o*^, 0^*o*^, +8^*o*^, +16^*o*^], and 3 vertical eccentricities, *y* = [−8^*o*^, 0^*o*^, +8^*o*^]. The yellow ellipses in each panel of the 3 *×* 5 grid of Fig. 11A depict Gaussian ellipsoids fitted to the retinal space–referred RF maps of synthetic mRGCs at the examined eccentricities. The small and non-systematic orientation biases in the retinal space–referred RF maps emerge due to the pooling of multiple cones by the RF center mechanism and are reminiscent of RGC mosaics mapped *in vitro* [32].

**Fig. 11.**
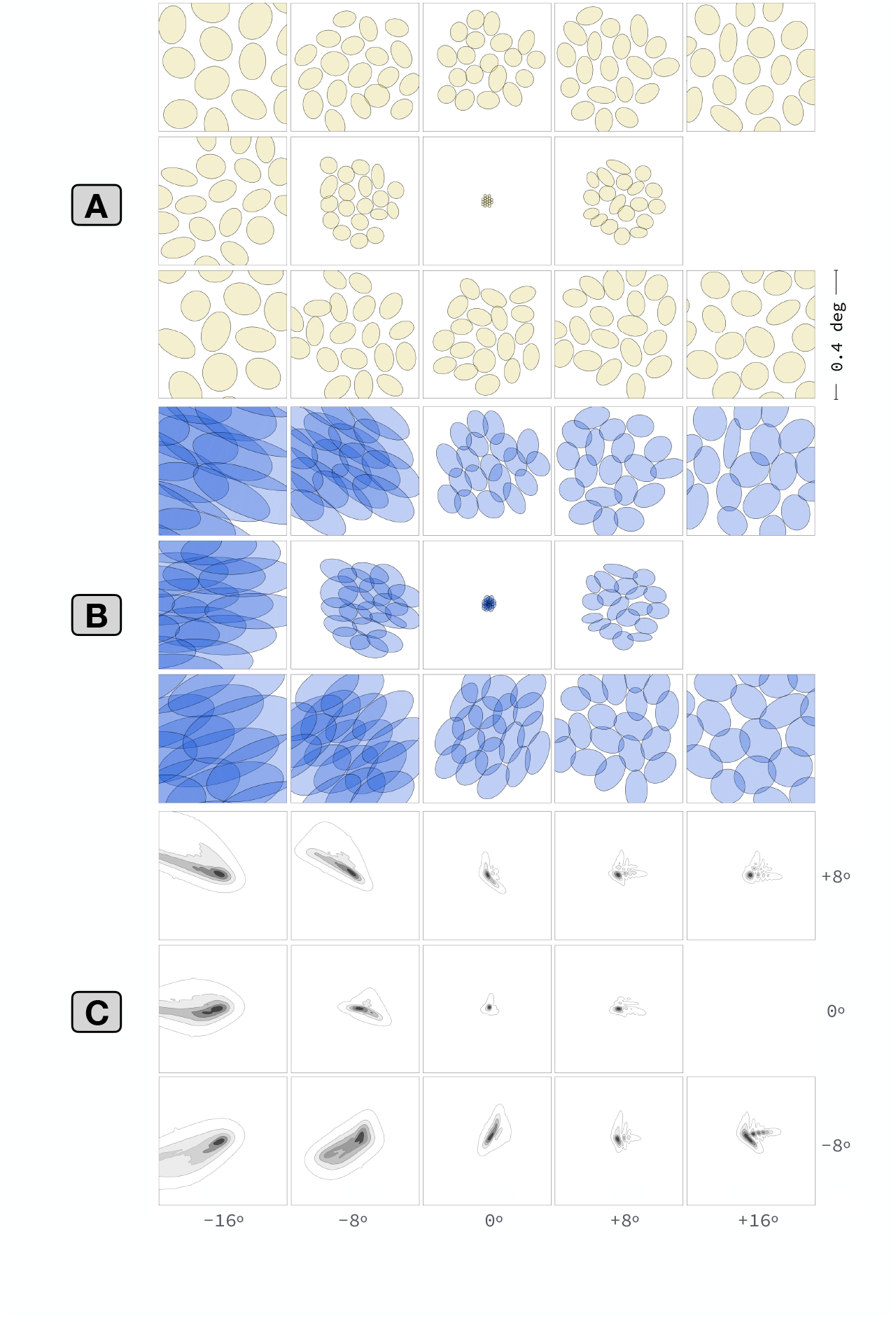
Retinal vs. visual space–referred mRGC RF maps across the retina. Illustration of the effect of physiological optics on visual space–referred spatial RF maps of synthetic mRGCs across eccentricity. **A:** Retinal space–referred spatial RF maps at different (x,y) eccentricities. Within each panel, yellow contours depict Gaussian ellipsoid fits to RF maps of up to 19 cells from a single mRGC mosaic. RF maps are computed using white noise stimulation under diffraction limited optics. **B:** Visual space–referred spatial RF maps of the same cells, computed under physiological optics appropriate for the cells’ eccentricity. **C:** Point spread functions of physiological optics at corresponding eccentricities.

The blue ellipses in Fig. 11B depict Gaussian ellipses fitted to the visual space– referred RF maps of the same cells. Note that there are striking and systematic orientation biases in these visual space–referred RF maps, which emerge due to the characteristics of physiological optics, whose point spread functions (PSFs) are depicted in Fig. 11C. Clearly, the shape of the PSFs, especially at peripheral locations is a major determinant of the visual space–referred RFs in mRGCs.

Overall, this analysis demonstrates that there can be substantial differences between *in vivo* and *in vitro* estimates of the spatial RFs of mRGCs, and again highlights the notion that inferences regarding retinal wiring from *in vivo* measurements must be evaluated in the context of the effect of the physiological optics. Indeed, in recent on-going work, [40], we have shown the importance of such analyses in assessing inferences regarding cone wiring to mRGC surrounds based on *in vivo* measurements of mRGC spatio-chromatic RFs.

### 3.5 Validity of the Difference of Gaussians model applied to *in vitro* responses of mRGCs in retrieving their spatial pooling characteristics

In our synthetic mRGCs, the spatial characteristics of cone pooling within the RF center and the RF surround *component* mechanisms are known by design. This allows us to test how well one can predict these characteristics from Difference of Gaussian model fits to *in vitro* measurements of mRGC STFs, where the RF center and surround mechanisms are driven simultaneously in the absence of optics [16]. Results of this analysis are illustrated in Fig. 12.

**Fig. 12.**
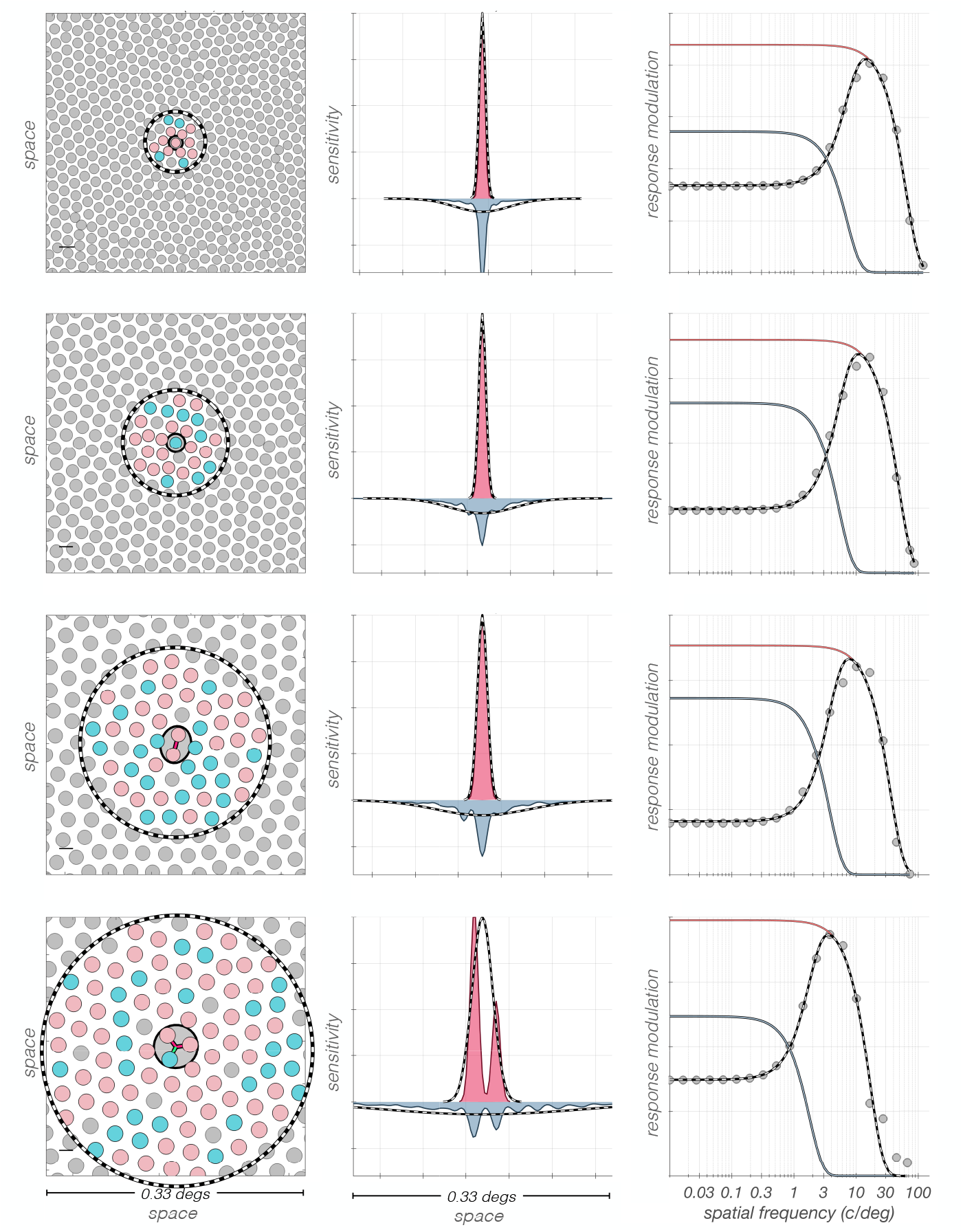
Spatial characteristics of mRGC surround cone pooling inferred from DoG model fits to retinal space–referred STFs are not accurate. The correspondence between actual and inferred surround cone pooling is examined for exemplar synthetic mRGCs at four eccentricities. **Left panels:** Cone input maps of exemplar synthetic mRGCs. **Middle panels:** Line weighting functions of cone inputs pooled by the RF center and surround mechanisms are depicted by pink and slate filled histograms, respectively. Dashed lines represent the line weighting functions inferred from DoG model fits to the model cells’ retinal space–referred STFs. **Right panels:** Retinal space–referred STFs, depicted by gray disks, are computed for stimuli delivered to the retina via diffraction limited optics. The DoG model fits to the computed STFs are depicted by the dashed black lines, with the corresponding center and surround component STFs depicted by the pink and slate lines, respectively.

The cone pooling maps of four exemplar mRGCs are depicted in the left column. The cells in the top two rows both have RF centers with a single cone input, whereas the cell in the third row has a 2-cone RF center, and the cell in the fourth row has a 3-cone RF center. The pink and maroon histograms depicted in the middle column of Fig. 12, are the y-axis integrated cone pooling weights within the RF centers and surrounds, respectively, and the superimposed dashed lines depict the center and surround line weighting profiles, as estimated by fitting the DoG model to the cells’ retinal space–referred STFs, which are depicted in the right column by the gray disks. Note that although the fitted DoG models (solid lines) captures the computed retinal space–referred STFs accurately for all cells, the inferred spatial RF profile, depicted by dashed lines in the middle column, do not accurately capture the surround mechanism cone pooling regions, which are depicted by the slate histograms. The discrepancy between actual and inferred surround pooling is most obvious in the two top cells which have single-cone RF centers, and becomes less pronounced as the RF center size increases. The discrepancy involves both the spatial extent and the peak sensitivity of the inferred surround pooling, which is estimated by the DoG model to be more diffuse with a weaker peak sensitivity than the cell’s actual surround cone pooling.

It is perhaps not surprising that the DoG model does not do a good job of fitting the model cell surrounds, given that they were constructed as double exponentials to match the spatial properties of H1 horizontal cells. The key point, however, is that the DoG model fits to the observable composite STFs are quite good. These observations suggest that caution should be exercised when inferring mRGC RF surround properties from DoG model fits to *in vitro* STF measurements.

### 3.6 Applications

We [2, 3, 6], and others [46–48] have reported on how the representation of visual information at the level of the cone mosaic shapes visual performance, in our case by exploiting the ISETBio image computable model of cone excitations. The transformation from cone excitations to RGC responses further shapes the information available for perceptual decisions, and we can interrogate our linear spatio-chromatic RF model of the ON-center mRGC mosaic to understand how the information available from this neuron class differs from that at the cone mosaic.

In this section, we present two example computations of this nature. Our goal is to illustrate how our model may be exploited in this way, and not to present a full analysis in either case. Even these initial calculations, however, provide interesting insight.

#### 3.6.1 Achromatic and chromatic spatial contrast sensitivity

We used a computational observer approach to compute spatial contrast sensitivity functions (CSFs) for achromatic and L-M cone opponent stimuli, based both on the representation at the cone mosaic and on the representation provided by our mRGC model. To do so, we computed responses to drifting gratings of varying spatial frequency, *ω*.

For the achromatic gratings, the L–, M– and S–cone contrast component gratings were in phase, *C*^*L*^(*ω, x, y*) = *C*^*M*^ (*ω, x, y*) = *C*^*S*^(*ω, x, y*). For the L − M gratings, the L– and M–cone contrast components were in antiphase, *C*^*L*^(*ω, x, y*) = −*C*^*M*^ (*ω, x, y*), and *C*^*S*^(*ω, x, y*) = 0. For all stimuli, mean (*xy*) chromaticity was (0.30, 0.32) and mean luminance was 100 *cd/m*^2^. Stimuli were simulated as presented on a typical CRT monitor, but with 20-bit channel DACs, to avoid intrusion of quantization effects.

For each eccentricity we studied, we oriented the gratings so that they were aligned with the axis of elongation of the optical point spread function at that eccentricity. Stimulus size was specified so that it extended over the area spanned by the input cone mosaic of the employed mRGC mosaic. The size of the mRGC mosaics was varied between eccentricities so as to achieve nearly equal numbers of mRGCs for mosaics between which we wished to compare performance.

Cone fundamentals vary with eccentricity because of variation in macular pigment density and photopigment axial density, and this variation is captured by ISETBio. Therefore, in these computations, stimuli were designed using cone fundamentals specific to the eccentricity of the employed mRGC mosaic.

At present, our mRGC model does not include spike generation or response noise. Therefore, in these computations we modeled response variability by adding zero mean Gaussian noise to the noise-free model mRGC responses. This approximation allows us to examine relative sensitivity across stimuli and eccentricity, but the overall level of predicted sensitivity is arbitrary. Given the choice of Gaussian noise, we used a template matching computational observer decision rule, with templates provided by the noise-free responses to the stimuli being discriminated. For comparing computational observer performance at the mRGCs with that at the cones, we also adopted the Gaussian noise approximation for the cone excitations, and used the template matching decision rule.

To estimate contrast sensitivity, we varied, for each tested spatial frequency, *ω*, the contrast of the test stimulus and identified threshold contrast, *C*_threshold_(*ω*), as that for which the probability of correctly identifying the test versus a zero contrast stimulus was 80.6%. Contrast sensitivity was defined as *CSF* (*ω*) = 1*/C*_threshold_(*ω*). Estimates of so computed contrast sensitivities at three eccentricities are depicted in Fig. 13. The left panels of Fig. 13 depict contrast sensitivities of mRGC mosaics and of their input cone mosaics for stimuli viewed through typical human optics. For comparison, the right panels depict corresponding calculations for stimulus viewed under diffraction-limited optics with no chromatic aberration, as might be measured using adaptive optics. The comparison between left and right panels helps understand which effects in the computed CSFs have their origin in the optics or sampling by the cone mosaic, and which should be attributed to retinal processing through to the mRGCs.

**Fig. 13.**
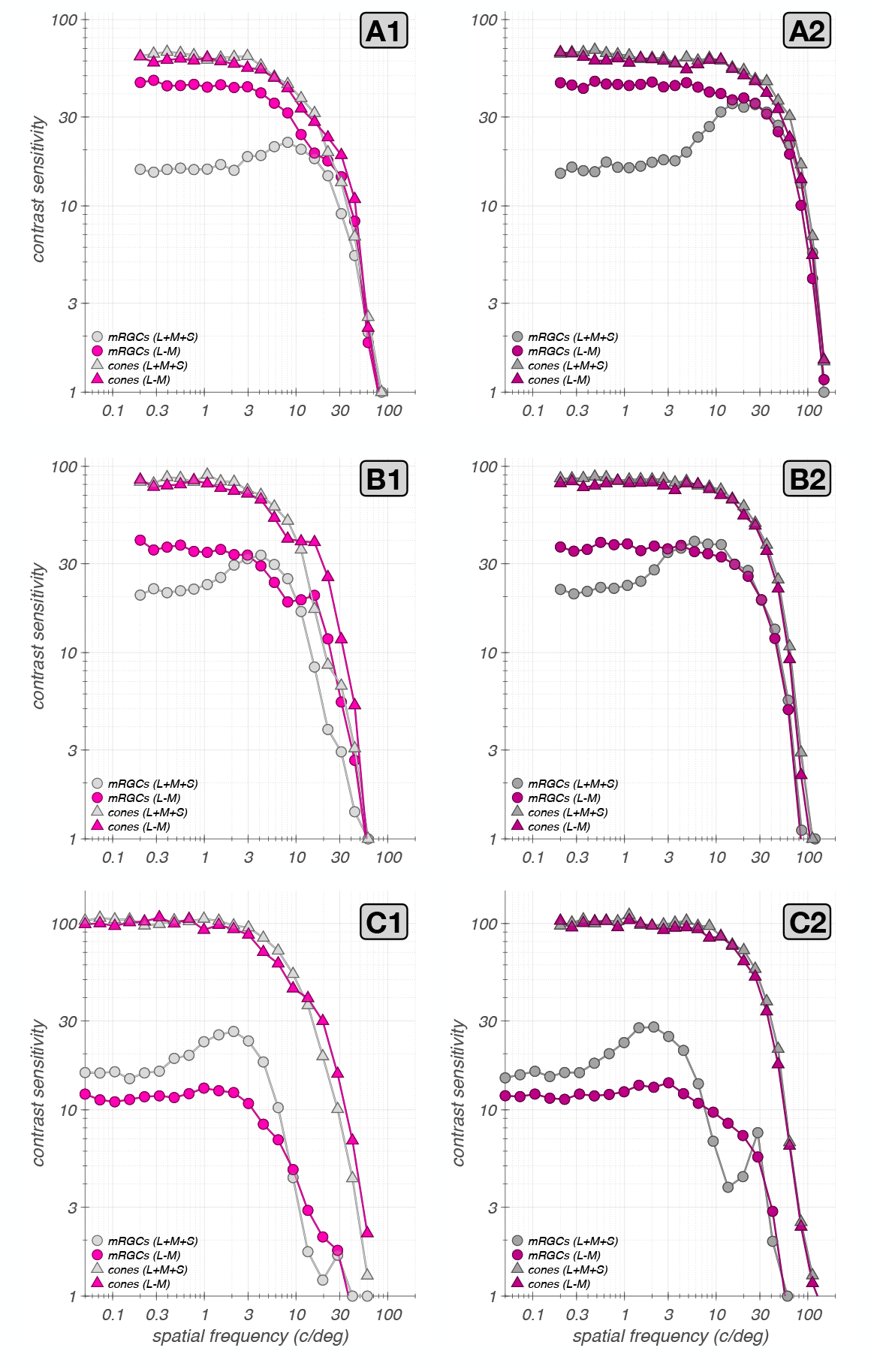
Computational observer spatial CSFs. Left column depicts CSFs computed with stimuli that are delivered to the retina through typical human optics. Right column depict CSFs computed with stimuli that are delivered to the retina under diffraction limited optics. **A1:** CSFs for a 0.6° ×0.6° foveal mRGC mosaic and of its input cone mosaic, depicted by disks and triangles, respectively. Gray: achromatic; pink: L-M. This mosaic contains 4628 mRGCs. **A2:** Diffraction-limited CSFs of the same foveal mRGC mosaic. **B1 & B2:** CSFs for a 2.1° × 2.1° parafoveal mRGC mosaic synthesized at an eccentricity of 4° along the temporal meridian. This mosaic contains 4633 mRGCs. **C1 &C2:** CSFs, respectively for a 4.1° × 4.1° peripheral mRGC mosaic synthesized at an eccentricity of 14° along the temporal meridian. This mosaic contains 2195 mRGCs.

At the fovea, the CSFs based on the cone excitations, depicted by triangles in Fig. 13A1, are low pass for both achromatic and L-M stimuli. This is expected because there is no spatial antagonism at the level of the photopigment excitations, and because we do not incorporate spatio-temporal coupling that arises because of interactions between fixational eye movements and post-receptoral temporal filtering [49, 50]. On the other hand, the achromatic CSF at the mRGC mosaic exhibits a mild low-spatial frequency attenuation, which is due to the spatial antagonism between mRGC RF centers and surrounds. Note that the low frequency attenuation appears weaker than what is observed under diffraction limited optics, depicted in Fig. 13A2. This occurs because physiological optical blur carves sensitivity at the high frequency regime, thereby reducing the apparent effect of the mRGC surrounds on the CSF. We observed a similar effect in foveal macaque mRGCs whose responses were measured under adaptive optics conditions [10].

The L − M opponent CSF of the mRGC mosaic lacks the low-frequency attenuation seen for achromatic modulations because in foveal mRGCs, L − M cone opponent stimuli do not induce substantial spatial antagonism between their single cone RF centers and their surrounds. These observations, which are consistent with what is known regarding the L − M chromatic contrast sensitivity of the mRGC pathway [16, 51], demonstrate that L − M sensitivity exceeds achromatic sensitivity at low spatial frequencies, consistent with the literature [52].

At high spatial frequencies there is little difference between computational observer sensitivity to achromatic and L − M modulations. This is not true of human observers, where sensitivity drops more rapidly as a function of spatial frequency for red-green isoluminant gratings than for achromatic gratings either with [51] or without typical optical blur [53]. Although our L − M opponent CSFs are not precisely equivalent to the red-green isoluminant CSFs measured in many human experiments, this is not the primary source of the difference between computational and human observers. Rather, it is known that compared to ideal observers, humans lose foveal information available from the cones more rapidly as a function of spatial frequency for red-green than than for achromatic gratings [48]. Our example calculation here suggests that this information loss should not be attributed to the linear receptive fields of the mRGCs. We believe this is because optical blur dominates computational observer performance at high spatial frequencies and the single cone RF centers of foveal mRGCs transmit information about each type of stimulus equally well; the surrounds have little effect at high spatial frequencies.

In the present calculations, the specific resolution limit, i.e., the spatial frequency at which sensitivity drops to 1, depends on the variance of the added Gaussian noise and is thus somewhat arbitrary. We have chosen a noise level that is low relative to human observers so that our computations show the behavior in the high-spatial frequency regime more fully than would psychophysics conducted through natural optics.

As we move to more peripheral locations, additional features of the CSF emerge. Figs. 13B1 and 13C1 depict results of computations at 4°. Note that under physiological optics viewing (Fig. 13B1) there is a spatial frequency regime in which L − M sensitivity exceeds the corresponding achromatic sensitivity, with the L − M CSF having a notched shape. We have reported this observation in conference abstract form [54]. It occurs because of the wavelength dependent defocus that is introduced by longitudinal chromatic aberration (LCA), which can change the spatial phase of the L– and M–cone stimulus components in the retinal image. Consistent with this interpretation, the notch is present in the CSFs both at the cones and at the mRGCs on the left, but not under diffraction-limited optics (Fig. 13B2), where LCA is zero. Similar effects have been observed for S-cone CSFs [55]. We have presented in abstract form experimental results that suggest that these effects occur in measurements of the human L − M spatial CSF [56].

Comparison of the cone–based CSFs in Fig. 13A1 with those in Fig. 13B1 and Fig. 13C1 also reveals the effect of stronger optical blur with eccentricity, which increases the rolloff of the CSFs with spatial frequency. Similar comparison of the mRGC-based CSFs shows additional rolloff introduced by the increasing size of mRGC RF centers with eccentricity.

Additional observations are notable at 14° (Figs. 13C1 and 13C2). First, a notch arises in the achromatic CSF at high spatial frequencies for the mRGC CSF that is not apparent in the cone CSF. This seems unlikely to be an optical effect, because it is more salient in Fig. 13C2 where optical effects are not present. Additional computations (not shown), indicate this notch is orientation dependent and has to do with the precise alignment of individual cones with the receptive field of an mRGC. We do not explore it further here. We do note that our computational observer is with respect to a noise level that makes it more sensitive than the human observer, so that the notch shown in Fig. 13C1 would be unlikely to be revealed with psychophysics conducted with natural optics. It is an interesting question as to whether it could be observed under adaptive optics conditions.

Second, the L − M advantage over the achromatic CSF is reversed at 14° of eccentricity. This is because at such high eccentricities, the L − M signal is reduced by the increased mixing of L– and M–cone signals within the larger mRGC RF centers and surrounds. Careful comparison of this effect with computational observer predictions for various choices of the model’s spatial homogeneity/spectral purity tradeoff parameter, *ϕ*, is an interesting future direction.

#### 3.6.2 Chromatic contrast sensitivity of synthetic mRGC mosaics: dependence on eccentricity

As a second example application, we examined chromatic sensitivity for uniform fields modulated in different directions in the LM–cone contrast plane. We used the same computational observer approach described above, and evaluated threshold for stimuli whose contrast was modulated in time. The cone contrasts of stimuli at different chromatic directions, *θ*, on the LM plane were:

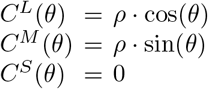

For each *θ*, we varied *ρ* to find its threshold value for discriminating that modulation direction from a zero contrast stimulus with a probability of 0.806. To summarize the computed thresholds across the different chromatic directions, we fit ellipses to the locus of threshold contrast points. Fig. 14 depicts computational observer thresholds for synthetic mRGC mosaics and for their input cone mosaics at different eccentricities.

**Fig. 14.**
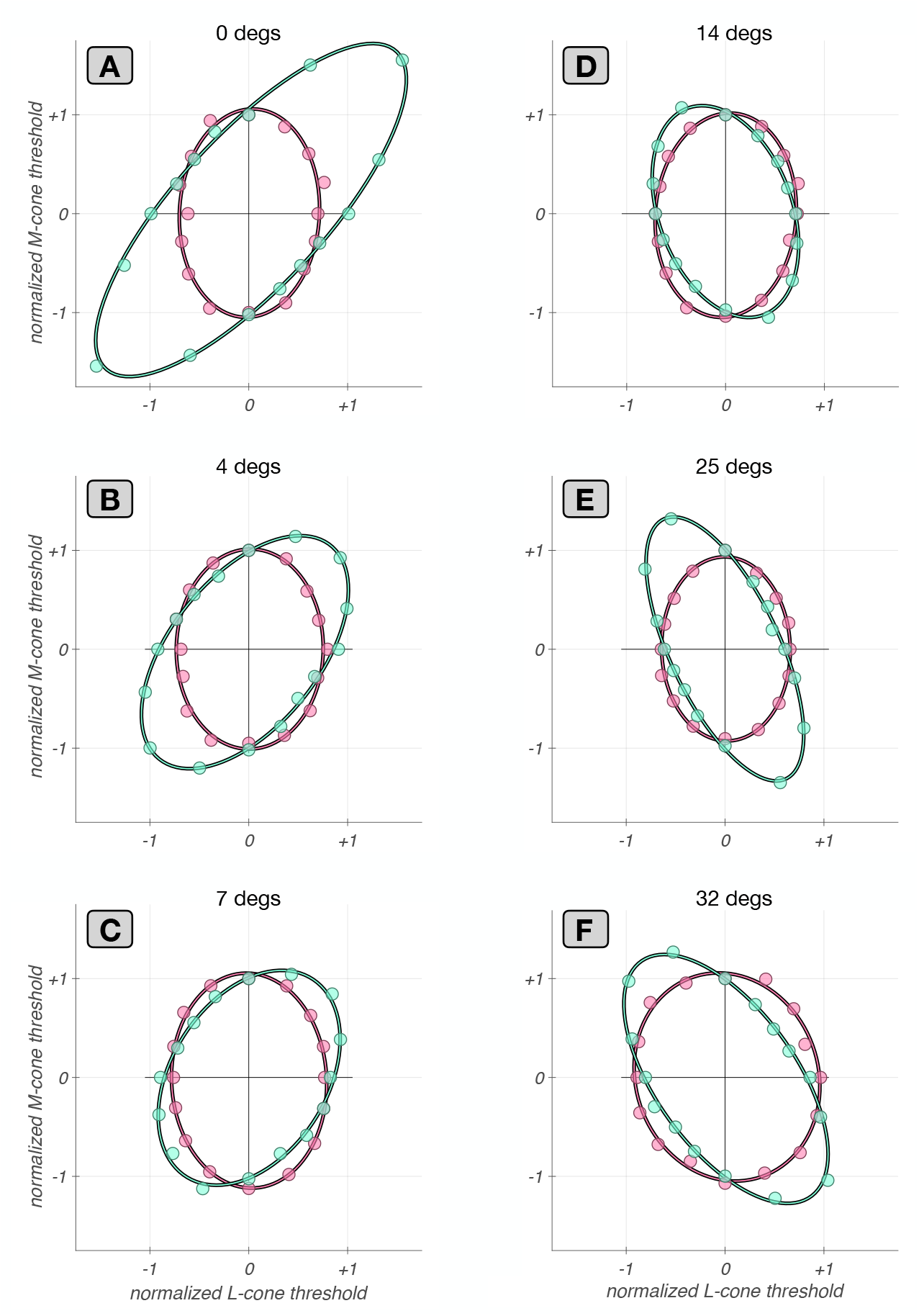
Chromatic contrast sensitivity of synthetic mRGC mosaics: dependence on eccentricity. Discrimination thresholds along the L/M-cone contrast plane of mRGC mosaics (green disks) and of their input cone mosaics (pink disks), computed for uniform field stimuli (0 c/deg). **A :** Data from a 0.6°× 0.6° foveal mRGC mosaic. **B :** Data from a 2.1° ×2.1° parafoveal mRGC mosaic synthesized at an eccentricity of 4° along the temporal meridian. **C :** Data from a 3.2° ×3.2° parafoveal mRGC mosaic synthesized at an eccentricity of 7° along the temporal meridian. **D :** Data from a 4.1° ×4.1° peripheral mRGC mosaic synthesized at an eccentricity of 14° along the temporal meridian. **E :** Data from a 6°× 6° peripheral mRGC mosaic synthesized at an eccentricity of 25° along the temporal meridian. **F :** Data from a 9° × 9° peripheral mRGC mosaic synthesized at an eccentricity of 32° along the temporal meridian.

How computational observer sensitivity changes with eccentricity depends on how stimulus size is covaried with eccentricity, as does human sensitivity (e.g. [57]). Comparison of the magnitude of sensitivity for cone- and mRGC-based computational observers depends on how the noise levels are chosen. For these example calculations, we focus on the shape rather than magnitude of the elliptical threshold contours. Each contour shown in Fig. 14 is normalized so that the threshold along the M cone direction is equal to one.

The first observation is that the normalized contours for the cone-based observer are similar across eccentricities and align with the L- and M-cone contrast axes. They are more elongated in the M-direction because our mosaics have more L cones than M cones. The alignment with the axes is expected [58], and the similarity of the normalized shapes occurs because this shape depends primarily on the relative numbers of L and M cones.

The contours for the mRGC-based computations, in contrast, change markedly with eccentricity. For the foveal mosaic, the threshold ellipse is highly elongated along 45° in the cone-contrast diagram, indicating that the highest discrimination thresholds occur when *c*^*L*^ = *c*^*M*^ and lowest thresholds occur when *c*^*L*^ = −*c*^*M*^ . This difference in comparison to the cone-based computations is a consequence of the chromatic-opponency of the mRGC RFs in the fovea. These have single cone centers, and thus opponency between the centers and the surrounds as the surrounds draw on mixed cone-types [59, 60]. The opponency leads to the observed contour orientation because it leads to cancellation of non-opponent L– and M–cone signals for low spatial frequency stimuli [58, 61].

As eccentricity increases, the contours first become less elongated and then elongation starts increasing again but along the 135° rather than the 45° axis. This is because the cone non-selective wiring model we implemented leads to progressively less opponency with increasing RF center size [16, 59, 60].

Although the qualitative features that emerge from this example calculation are understood in the literature, the example illustrates that our model enables this type of calculation to be made quantitatively in a way that takes chromatic aberration, stimulus size and spatial frequency, and retinal position into account. Of particular interest to us will be exploring how this type of threshold contour varies with the center wiring parameter *ϕ* of our model, which specifies the tradeoff between spatial homogeneity and spectral purity of mRGC RF centers.

## 4 Discussion

We developed an image computable model of the linear spatio-chromatic RF mosaic of mRGCs across the retina. The model extends our image-computable cone mosaic model [2, 3] by adding a layer of mRGCs which pool signals directly from the cone mosaic. The connectivity between cones and mRGCs is derived using a simulation framework that integrates anatomical, physiological and optical quality data, all of which vary across eccentricity.

By explicitly modeling the optics and photoreceptors, rather than directly expressing the RFs in terms of the stimulus, we are able to link our model with both *in-vitro* and *in-vivo* data, and to make predictions over a range of experimental conditions that are otherwise difficult to compare. These include psychophysical and physiological measurements made through physiological optics (natural viewing conditions), interferometric and adaptive optics techniques that bypass or correct for optical aberrations, and *in-vitro* physiology, where the natural optics are not present.

To build the model we had to overcome the challenge that current data about mRGC properties are incomplete and, where they exist, may come from different species, different measurement modalities, and from different eccentricities. For example, there are *in-vivo* measurements of mRGC linear receptive fields across the retina [17], but physiological optics blur the stimuli so that they do not constrain mRGC input at the cone-by-cone resolution we seek. On the other hand, although there is single cone-resolution connectivity data from *in-vitro* physiology [15], these data are currently limited to large eccentricities (≥ 25^*o*^). Thus, we developed a modeling framework that allows integration of data from multiple sources. This framework is an important contribution in its own right; we expect it will be useful to us and others, for incorporating new data that become available and for modeling other RGC classes.

We showed that the model captures visual space–referred spatial RF properties of macaque mRGCs recorded *in-vivo* across eccentricities, as well as retinal space– referred spatial RF properties of macaque mRGCs recorded *in-vitro*. We also showed that physiological optics plays a major role in shaping the visual space–referred spatial RF properties, so that inferences regarding retinal circuitry made from *in-vivo* measurements need to be evaluated in the context of the optics. Further, we showed that even under *in-vitro* conditions, where the optics are eliminated, the traditional approach of fitting a Difference of Gaussian model to spatial responses can lead to incorrect assessments of the properties of cone pooling in the mRGC surrounds.

### 4.1 Applications

We employed an early version of the current model to interpret measurements of foveal macaque mRGCs measured *in-vivo* using adaptive optics [10]. Specifically, the model allowed us to relate the adaptive optics measurements to *in-vivo* measurements conducted under physiological optics. For this purpose, the ability to move back and forth between retinal and visual space-referred representations was critical.

We are currently employing the model to assess inferences regarding the wiring of cone inputs to mRGC RF surrounds based on spatial RF measurements conducted *in-vivo* [19]. Specifically, we are analyzing the substantial effect that chromatic aberration plays in shaping mRGC responses to cone isolating stimuli, and how these effects can help reconcile tension between results from *in-vivo* physiology on the one hand and results from anatomy and *in-vitro* physiology on the other [40].

In parallel on-going work, we deploy the model to understand how the spatiochromatic properties of the ON-center mRGC mosaic influence the information available for human spatio-chromatic vision, by applying computational observer analyses to the mRGC representation we compute [54, 56]. Although additional model components will influence this representation, for threshold tasks where the stimulus perturbations are small, we expect the linear approximation to hold sufficiently well that the results will be informative.

In this work, we presented examples of this type of computation, to illustrate how the representation at the mRGCs differs from that at the cone mosaic, and how this varies with eccentricity.

### 4.2 Limitations and Future Directions

#### 4.2.1 Human versus macaque

When available, we used human data to guide model development, in order to maximize the usefulness of the model in predicting human performance. Even if this had not been our goal, we would have had to bring in human data to characterize the physiological optics across the visual field, as such data are not currently available in macaque.

At the same time, not all the required data are available for human: although measurements of cone and mRGC density and physiological optics across the retina are available, physiological characterizations come from the macaque.

The need to mix data across the two closely related species produces tension in cases where the parameters for the two species differ. An example is the different cone densities in the far periphery [45], which intrudes on the interpretation of the comparison between our model and *in-vitro* physiology in that retinal region. As more data become available in both species, and as species differences come more fully into focus [62], our approach should allow more fully differentiated models to be developed targeted at each.

#### 4.2.2 Noise, nonlinearities and temporal dynamics

Although the current model captures fundamental aspects of the visual representation at the level of the mosaic of ON mRGCs, there are known characteristics of mRGCs that it does not account for. These include static and spatial nonlinearities, temporal filtering, spike generation, and physiologically constrained response noise. The modeling framework we developed is extensible however, so that these components may be included through future work.

Response variability models are available for macaque mRGCs, as descriptions of spike generation mechanisms [26, 63, 64]. In addition, we can incorporate nonlinearities, such as (a) adaptation effects introduced through the phototransduction cascade [65], (b) compressive and expansive static nonlinearities in the output of mRGCs [23, 64], and (c) spatial nonlinearities introduced by rectifying sub-units within the RFs of mRGCs [21, 22]. Explicit inclusion of photocurrent-based responses in the input to the mRGCs introduces a temporal component to the response model [65]. In addition, a second temporal filter may be added, such that when combined with the photocurrent filter will yield the bandpass filter characteristics observed in macaque mRGCs [25].

Our current model does not represent explicitly the properties of the retinal circuitry (horizontal, bipolar, and amacrine cells) that produces the mRGC response properties, as we have opted instead to work towards a functional model that describes those properties. A complementary mRGC modeling approach that does consider some of these cell types explicitly has recently been published [66]. We note however, that some of the processing performed by these other retinal cell types is incorporated implicitly in the current cone-to-mRGC model, such as the parametric form of the surrounds inherited from H1 cells. The framework we developed is designed so that it would be possible to interpose explicit models of intermediate retinal cell types. Representing the action of different cell types explicitly may in the longer run be an effective way to account for response nonlinearities in the mRGCs, or in other classes of retinal ganglion cells. Moreover, using our framework to model other cell classes may be of interest to those seeking to interpret responses of those classes *per se*, or in the retinal mechanisms that produce RGC response properties.

#### 4.2.3 OFF mRGC mosaic

Because we model the linear RF, the distinction between ON and OFF mRGCs is subtle. However, our model should be thought of as a model of only the ON mRGCs because the synthetic cells only pool signals from L– and M–cones. This is believed true for ON mRGCs, but recent evidence suggests that OFF mRGCs draw upon all three types of cones in their RF centers [15, 30, 31]. Incorporating S-cone input into an OFF–center mRGC model is straightforward.

Another question that arises when considering a model of OFF mRGC mosaic is how to split the density of mRGCs in two populations at different eccentricities. In the current model, the ON mRGC density was assumed to be half of all mRGCs across all eccentricities. This seems reasonable for central retina where mRGC centers draw primarily on a single cone and where anatomical evidence suggests that each cone provides input to the center of one ON and one OFF midget bipolar cell. However, there is evidence that the RFs of peripheral ON midget (and parasol) RGCs are larger than their OFF counterparts in both human and macaque retinas [33]. This implies that the density of ON RGC cells might be lower in the periphery than that of OFF cells, given that ON and OFF mRGCs have similar RF overlap [32]. One idea is to treat the asymmetry between ON and OFF mRGC RF densities in an eccentricity-dependent manner, similar to the way we encoded a variable-with-eccentricity RF center overlap.

Finally, when adding an OFF mRGC mosaic one should allow for the possibility of coordination between the ON and the OFF submosaics, to account for recent observations regarding systematic shifts in the spatial layouts of ON and OFF mRGCs [67].

## Using the software

The developed software for synthesizing ON mRGCm mosaics across the retina and for computing with them is part of ISETbio and is freely available at https://github.com/isetbio/isetbio. An introduction to using the mRGCmosaic software is available at: https://github.com/isetbio/isetbio/wiki/Retinal-ganglion-cell-(RGC)-mosaics, and a number of MATLAB tutorials specific to the mRGCmosaic can be found at https://github.com/isetbio/isetbio/tree/main/tutorials/mrgc.

These tutorials demonstrate (a) how to use mosaics of ON mRGCs that have been synthesized at a number of eccentricities, and (b) how to build and validate mRGC mosaics at any desired eccentricity, using a number of design choices. A summary of current available tutorials is shown in Table 1.

**Table 1.** List of tutorials for computing with mRGC mosaics and de novo synthesis of mRGC mosaics.

| Tutorial name | Scope |
| --- | --- |
| *Computing with mRGC mosaics* |  |
| `t_mRGCmosaicVisualizeWithOptics.m` | Visualizes a previously synthesized mRGC mosaic and the optics that were used for its synthesis |
| `t_mRGCmosaicInspect.m` | Visualizes an mRGCmosaic and cone pooling maps of individual cells |
| `t_mRGCmosaicBasicComputation.m` | Perform a basic computation with an mRGC mosaic |
| *Synthesizing mRGC mosaics* |  |
| `t_mRGCmosaicSynthesizeAtStage1.m` | Denovo synthesis of the spatial position lattices of cones and mRGC RF centers (stage 1) |
| `t_mRGCmosaicSynthesizeAtStage2.m` | Denovo synthesis of an mRGC mosaic at different sub-stages of cone-to-mRGC RF center connectivity (stage 2) |
| `t_mRGCmosaicSynthesizeAtStage3.m` | Denovo synthesis of an mRGC mosaic at different sub-stages of cone-to-mRGC RF surround connectivity (stage 3) |

## Acknowledgements

We thank Lisa Croner and the late Ehud Kaplan for help with procedures followed during collection of the macaque *in vivo* data, Greg Field for providing insight on details regarding *in vitro* macaque data, and E. J. Chichilnisky, Tyler Godat, David Williams, and Fangfang Hong for helpful discussions.

## Declarations

### Ethics approval and consent to participate

Not applicable

### Consent for publication

Not applicable

### Data availability

Datasets (mRGCmosaics) generated during the current study are available at: https://github.com/isetbio/isetbio/tree/main

### Materials availability

Not applicable

### Code availability

The code used to generate the data, and various tutorials on how to use the software are available at: https://github.com/isetbio/isetbio/tree/main

An introduction to using the software is available at: https://github.com/isetbio/isetbio/wiki/Retinal-ganglion-cell-(RGC)-mosaics

### Author contribution

NPC: conceptualization, algorithm development, coding, data curation, validation, visualization, writing of original draft

DHB: conceptualization, coding, reviewing and editing of manuscript

BW: conceptualization, coding, reviewing and editing of manuscript

## Appendix A Deriving cone weights to the mRGC RF centers

### A.1 Local topology–based convergent connections (stage 2A)

During the first sub-stage of cone-mRGC RF center connectivity, cones are connected to single mRGC RF centers based on the local topology of the 2 lattices. Starting with the cell whose RF center is at most central location of the mRGC lattice, we connect *n*_p*ool*_(*ϵ*) number of L– and M–cones to it,

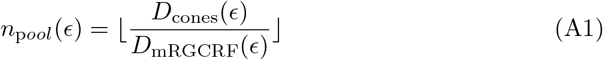

where *D*_cones_(*ϵ*) and *D*_mRGCRF_(*ϵ*) are the local spatial densities of the cone mosaic and of the mRGC RF centers, respectively, at the eccentricity, *ϵ*, of the target mRGC. We draw from the nearest *n*_p*ool*_(*ϵ*), typically 6, cones that have not yet been connected and whose distance to the mRGC RF center does not exceed a fraction of the local mRGC RF center spacing. This fraction is a parameter of the model and for the work presented here was set to 0.6.

Continuing with these assignments of cones to mRGC RF centers, we move outward to more peripheral locations in the mRGC RF mosaic, connecting cones to each mRGC center. Any L– and M–cones that remain unconnected at the end of this sub-stage are then connected to their nearest mRGC center, so that all cones are connected to one mRGC center.

This sub-stage can result in local inhomogeneities in both the number of cones and the type of cones pooled within neighboring RF centers. These inhomonegenities are smoothed out as part of the next sub-stage.

### A.2 Optimizing cone connections to mRGC RF centers (stage 2B)

In the second sub-stage of the RF center connectivity, convergent connections from multiple cones to single mRGC RF centers are optimized according to a desired balance between spatial and spectral inhomogeneities. This is achieved by reassigning cones between nearby mRGC RF centers, which itself occurs in two steps.

In the first step, we allow cone reassignments to a target mRGC from neighboring mRGCs that have a higher input cone numerosity in their RF centers. In the second step, we allow cone swaps between a target mRGC and its neighbors, independently of their input cone numerosities.

The heuristics followed in the first step are as follows. We begin by targeting mRGCs with a single input cone and continue to target mRGCs with progressively higher input cone numerosity. Within each set of targeted input cone numerosity, mRGCs are sorted based on ascending retinal eccentricity. For each targeted mRGC we determine up to 6 neighboring mRGCs which have input numerosity that exceeds that of the target mRGC by at least 2 cones.

Cone reassignments from the candidate donor mRGCs to the target mRGC are executed in multiple passes. Starting with the neighboring mRGC of the highest input numerosity, we determine the best transfer of a single cone. If there are no eligible donor nearby mRGCs, we move to the next targeted mRGC. If there is a single candidate, we accept it and execute the cone transfer. If there are more than one candidates, for each candidate donor mRGC (up to 6) we compute a cost function, **C**, for reassigning each of its cones to the target mRGC, and pick the transfer that minimizes **C** across all cones and all candidate donor mRGCs. The cost function is described in more detail below.

Once the optimal cone transfers for each mRGC of the targeted input cone numerosity are executed, we move to the next pass, examining possible transfers from neighboring mRGCs of one lower input cone numerosity than before, but still higher than the input cone numerosity of the targeted mRGCs. Once all passes are executed, this process is repeated, now targeting mRGCs with increasing input cone numerosity, until all input cone numerosities have been targeted.

In the second step, we only allow for cone swaps between an mRGC RF center and one of its neighbors. For each mRGC of the targeted input cone numerosity, we determine its 6 closest neighbors, but now without regard to their input cone numerosity. For each of these neighboring mRGCs, we evaluate the cost function, **C**, for all possible combinations of cones from the target mRGC and cones from the neighboring mRGC and pick the combination that minimizes **C**. The selected cone swap is executed only if the post-swap value of **C** is lower than its pre-swap value. Multiple passes through the entire mRGC mosaic, are executed, with each pass targeting mRGCs with progressively higher input cone numerosity.

The cost function, **C**, employed to determine the optimal transfer/swap is based on the position and types of the cones pooled by the target mRGC, *t*, and the examined neighboring mRGC, *t*_*i*_. For each examined pair of mRGCs, 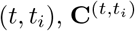 is defined as:

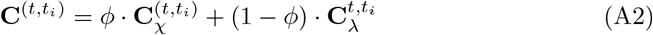

where 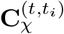 is a spatial compactness cost component, 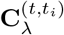 is a spectral purity cost component, and *ϕ* is a free parameter that controls the desired trade-off between spatial compactness and spectral purity of the RF centers.

The spatial compactness cost component, 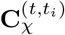, is defined as:

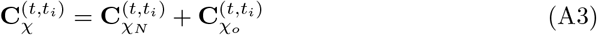

In Eq. A3, 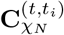 quantifies the differential input cone numerosity between the examined pair of mRGCs, and is defined as:

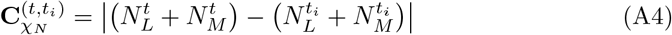

with 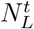 and 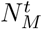 are the numbers of L– and M–cones pooled by the RF center of mRGC *t*, respectively.

The 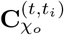 component of Eq. A3 is a spatial overlap cost component, defined as the inverse of the distance between the centroids, 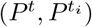, of the cones pooled by the two mRGCs, normalized by the sum of their spatial standard deviations, 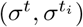:

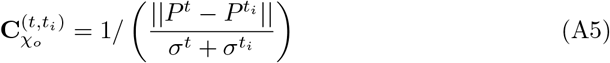

A low value of 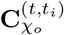 indicates low overlap between the sets of cones pooled by the examined pair of mRGCs and conversely, a high value indicates a large overlap.

The 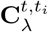 term, in Eq. A2, is a cost component quantifying the sum of spectral impurities of the pair of analyzed mRGCs:

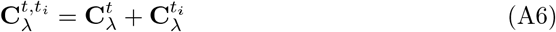

The spectral impurity, 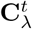, quantifies the degree of non-specificity, with regard to the type of cone, in the pooling withing the RF center of an mRGC, and is defined as:

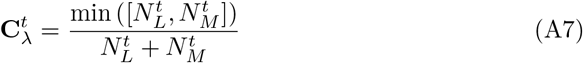

Values of 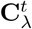 near zero indicate a low amount of mixture of L– and M–cones, and therefore a RF with a high degree of spectral purity, and conversely, values of 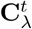, near 0.5, indicate an equal mixture of L– and M–cones, and therefore a RF center with a low degree of spectral purity.

The tradeoff parameter *ϕ* factor in Eq. A2, controls how the optimal cone transfer/swap is determined. When *ϕ* = 1, cone reassignments/swaps are selected so as to minimize the spatial compactness cost, when *ϕ* = 0, cone reassignments are chosen so as to minimize the spectral purity cost, and for intermediate values of *ϕ*, cone reassignments are chosen so as to minimize a ratio of the two costs.

### A.3 Divergent cone connections to multiple mRGC RF centers (stage 2C)

In the final sub-stage of establishing the RF center connectivity, the exclusivity of connections is relaxed, and cone connections are allowed to diverge to more than one mRGC RF center. This divergence is guided by *in-vitro* measurements of mRGC RF center overlap in the macaque [32].

According to these observations, neighboring mRGC RF centers abut at approximately one standard deviation of their Gaussian RF profile. One caveat of using these *in-vitro* measurements to establish cone divergence in the model, is that these measurements are only available in the far periphery (30–40 degrees), with no data available for more central locations. Anatomical studies suggest, however, that, in the central retina, there must be little to no divergence of cone signals to mRGCs RF centers, so we chose to implement an eccentricity-varying divergence in our model.

We begin by fitting an ellipsoid to the spatial pooling map of cones that are exclusively connected to the RF center of an mRGC, and extracting the rotation, *α*, and the major/minor axes, *σ*_*x*_, *σ*_*y*_ of the fitted ellipsoid. Next, a super-Gaussian ellipsoid, *G*(x, y, n), defined as:

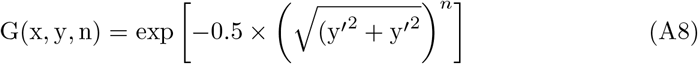

where:

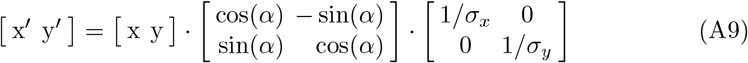

is computed by scaling the values of *σ*_*x*_, *σ*_*y*_ by a common factor, so that the value of G(x, y, n), evaluated at the most remote exclusively-connected cone(s) is *k × e*^−1*/*2^. The value of *k* is determined empirically so that RF maps of nearby mRGCs computed under diffraction-limited optics abut when their sensitivities drop to *e*^−1*/*2^ (per [32]). By varying the exponent of the super Gaussian, *n*, we model varying degrees of cone divergence. When *n* = 10, we obtain a flat-top Gaussian with very sharp fall-offs, modeling minimal cone divergence. When *n* = 2, we get a standard Gaussian modeling cone divergence that is consistent with the *in-vitro* measurements of RF center overlap at peripheral locations.

By varying *n* with eccentricity using a sigmoidal function we obtain a gradual transition in cone divergence with eccentricity. The slope and mid-point of the sigmoidal variation of *n* are currently chosen arbitrarily, with the only restrictions that above 15^*o*^, *n* is stable at 2.0, and below 7^*o*^, *n* is stable at 10.0. The weights of divergent cone-mRGC RF center connections are computed by evaluating the super-Gaussian ellipsoid at the positions of all cones in the vicinity of the examined mRGC.

## Appendix B Deriving cone weights to the mRGC RF surrounds

### B.1 Choosing physiology-based constraints for deriving surround cone weights in stage 3B

The optimization of the parameters of the surround cone pooling functions at each iteration is driven by the residual between the visual STF that is computed based on the surround pooling weights at the previous iteration and the Difference of Gaussians model fit to it, DOG(*ω*), which is given by:

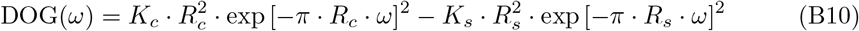

This aspect of the optimization captures the observation that DoG models provide a reasonable fit to *in-vivo* mRGC RFs. To ensure adherence to the *in-vivo* data of Croner & Kaplan, the DOG(*ω*) model fit is constrained so that the ratio of surround to center radii, *R*_*s*_*/R*_*c*_, and the ratio of surround to center integrated sensitivities, *K*_*s*_*/K*_*c*_ *×* (*R*_*s*_*/R*_*c*_)^2^, both remain within a specified tolerance value, *τ* = 0.15, from the mean values of the corresponding ratios for the Croner & Kaplan population of macaque mRGCs at the eccentricity of the synthesized mRGC.

Specifically, for the model’s *R*_*s*_*/R*_*c*_ ratio, we enforce

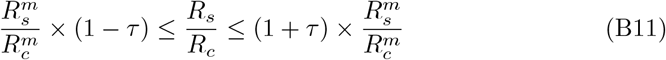

where 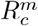 and 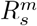 are the mean values of center and surround radii across the Croner & Kaplan population of macaque mRGCs at the eccentricity of the synthesized mRGC. The model’s *K*_*s*_*/K*_*c*_ *×* (*R*_*s*_*/R*_*c*_)^2^ ratio is constrained in the same way.

The residual between the visual STF and the Difference of Gaussians model fit to it, drives the optimization of the surround pooling function. This function is a double exponent (following the H1 horizontal cell spatial RF in the macaque [36]):

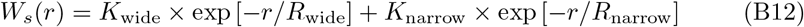

To ensure that the surround pooling function remains consistent with parameter values observed in macaque H1 cell [36], the optimization of *W*_*s*_(*r*) is constrained so that ratio of radii, *R*_narrow_*/R*_wide_, and the ratio of volumes, *V*_narrow_*/V*_wide_ = *K*_narrow_*/K*_wide_ *×* (*R*_narrow_*/R*_wide_)^2^, of the two exponentials both remain within a specified tolerance range of the macaque data.

In the present work, the tolerance range for *R*_narrow_*/R*_wide_ was set to [0.07, 0.35] for all mosaics, whereas the tolerance range for *V*_narrow_*/V*_wide_ was set to [0.01, 0.6] for mosaics at eccentricities ≤ 15°, to [0.3, 0.9] for eccentricities in 15° … 25°, and to [0.6, 1.3], for eccentricities ≥ 25°.

The joint manipulation of the tolerance values applied to the parameters of the DOG model fit to the vSTF, and to the parameters of the double exponential surround pooling model, *W*_*s*_(*r*), allows for different options for deriving spatial pooling functions in synthetic mRGC surrounds.

One option is to set very strict tolerances on the parameters of DoG model fit to the achieved vSTF, while allowing for a large tolerance in the parameters of *W*_*s*_(*r*). Results of this choice are depicted in the left-most column of Figure B1. A second option would be to allow medium tolerance levels in both the DoG model fit and the *W*_*s*_(*r*). Results of this choice are depicted in the middle column of Figure B1. A third option would be to enforce strict tolerances in *W*_*s*_(*r*), for example matching parameters of individual H1 horizontal cells, while allowing for a loose tolerance in the DoG model fit. Results of this choice are depicted in the right column of Figure B1. In the present work, we chose the second option.

**Fig. B1.**
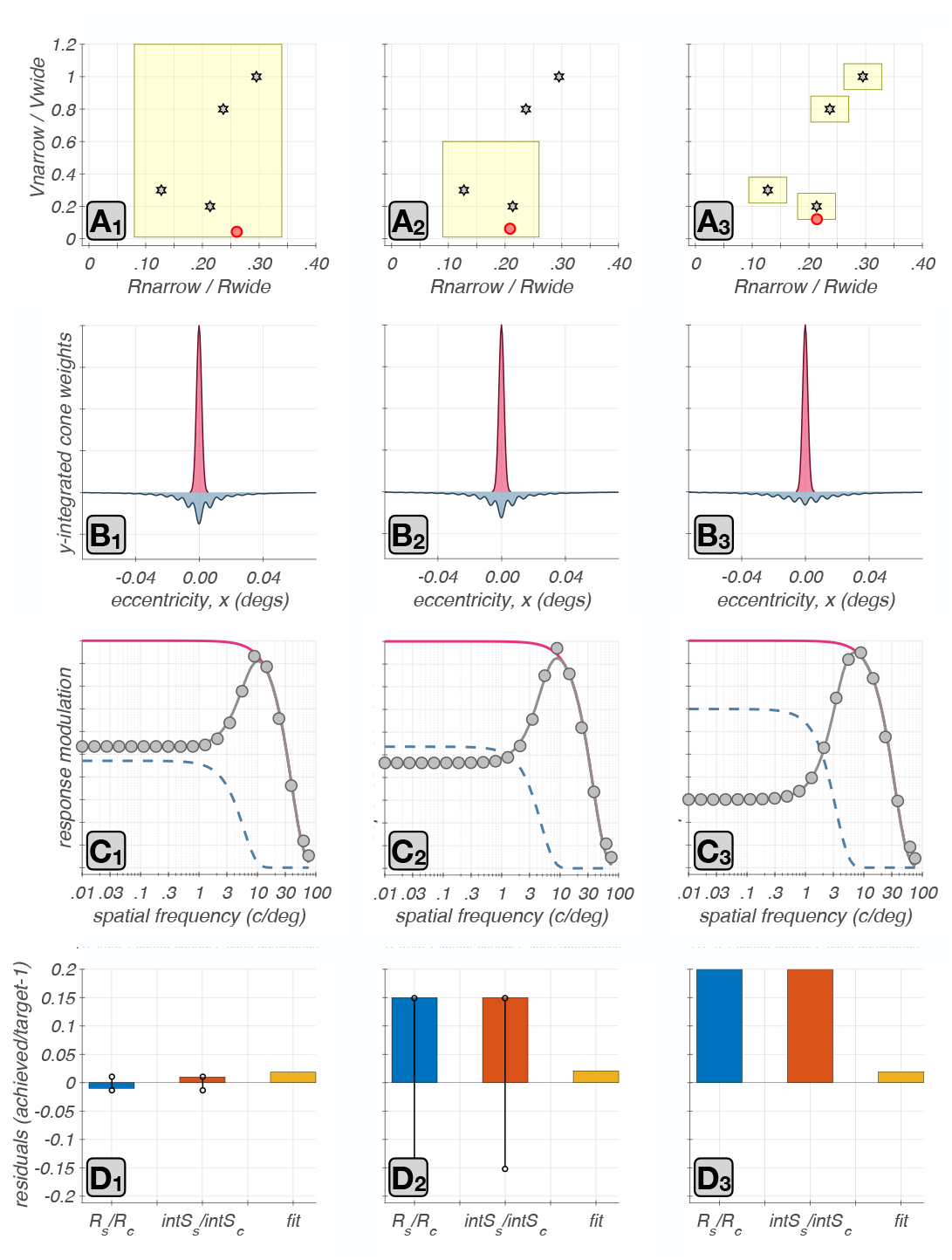
Effect of constraints on surround cone pooling. Results from three options for the constraints are depicted. In the left column, we see results from a tight tolerance in the parameters of the DoG model fit to the vSTF and a loose tolerance in the parameters of the double exponential surround pooling model, *W*_*s*_(*r*). In the middle column, we see results from medium tolerance in both sets of parameters. In the right column, we see results from a loose tolerance in the DoG parameters and a tight tolerance in the *W*_*s*_(*r*) parameters. **A1-A3 :** The tolerance range enforced during model fitting in the joint space of the two surround cone pooling related parameters, *V*_narrow_*/V*_wide_ and *R*_narrow_*/R*_wide_ for the three examined strategies is depicted by the yellow rectangles. Stars depict the corresponding parameter values in four macaque H1 horizontal cells from the study of Packer & Dacey. The red disk depicts the achieved parameter values under each strategy for an example foveal synthetic mRGC. **B1-B3 :** Line weighting functions of the retinal space referred center and surround cone pooling weights under the three examined strategies. **C1-C3 :** vSTF computed under the three strategies are depicted by the. disks, and the DOG model fit by the gray lines, with the red and blue lines depicting the center and surround components of the DOG model fit. **D1-D3 :** Blue and orange bars depict the residuals for the ratios of visual space - referred *R*_*s*_*/Rc* and *K*_*s*_*/K*_*c*_ × (*R*_*s*_*/R*_*c*_)^2^ ratios. Black circles connected by a black line depict the enforced tolerance range in these ratios. The enforced tolerance value in D3 was *τ* = 0.5, and is not visualized. Yellow bars depict the ||vSTF(*ω*) − DOG(*ω*)|| residual.

